# A statistical framework for identifying candidate genetic interactions from adaptive laboratory evolution data

**DOI:** 10.64898/2026.11.12.766281

**Authors:** Binhuan Sun, Adam M. Feist, Patrick V. Phaneuf, Alberto Santos

## Abstract

**Background:** Adaptive laboratory evolution (ALE) is widely used to improve microbial fitness under defined selective pressures. Mutations accumulated during ALE may interact epistatically, with one mutation’s effect depending on another. Understanding these interactions can help explain adaptive phenotypes and inform combinatorial strain engineering, but experimentally testing all mutation combinations is impractical. We developed a computational framework that uses gene-pair co-occurrence and mutual exclusivity patterns to identify candidates for experimental investigation of epistasis.

**Results:** We applied the framework to a curated compendium of 141 *Escherichia coli* ALE experiments from ALEdb, using Fisher’s exact test and DISCOVER as complementary statistical screens. Testing and multiple-testing correction were performed separately within each experiment, method, and direction (co-occurrence and mutual exclusivity). At a 5% false discovery rate threshold, Fisher’s exact test identified 74 significant gene pairs, comprising 36 co-occurrences and 38 mutual exclusivities. DISCOVER identified 5 significant pairs, all for mutual exclusivity and also significant under Fisher’s exact test. These shared pairs suggested alternative adaptive routes involving temperature tolerance, glycerol utilization, and association between RNA polymerase subunit genes. The two methods produced similar expected overlaps for Fisher-significant pairs, but DISCOVER’s broader null distributions yielded larger raw *p*-values. Of the 69 pairs significant only under Fisher’s exact test, 60 arose in experiments with multiple strain labels or involved a gene with only one distinct mutation recorded across all affected endpoints (final samples from lineages evolved in parallel), highlighting the need to examine genetic backgrounds and mutation origins when interpreting the results. Power analysis indicated that Fisher’s exact test required 70 independent endpoints for co-occurrence and 215 for mutual exclusivity to achieve 80% power under pre-specified scenarios. We implemented the framework as a reusable Nextflow pipeline that automates statistical screening of ALEdb and user-supplied mutation datasets.

**Conclusions:** By combining gene-pair screening with mutation-level interpretation, this reproducible framework narrows the search space for experimental investigation of epistasis and can contribute to combinatorial strain design. The power analysis provides sample-size benchmarks for planning future ALE experiments with adequate power to screen for candidate genetic interactions.

## Background

Adaptive laboratory evolution (ALE) is widely used to investigate microbial adaptation under defined selective pressures [1–4]. It involves propagating microbial populations over successive generations, and mutations that confer a fitness advantage can accumulate during this process. Genome sequencing of evolved lineages provides mutation datasets for investigating the genetic basis of adaptation. However, mutations observed in ALE experiments can include both neutral mutations and beneficial mutations with varying fitness effects [5]. Data-driven analyses have therefore been developed to identify and prioritize potentially beneficial mutations [6–8]. These candidate mutations can subsequently be reconstructed in a defined genetic background and experimentally evaluated for their phenotypic effects.

Combining beneficial mutations provides a potential route to greater improvements in fitness or other phenotypes of interest. However, their joint effect is not always predictable from their individual effects, because the effect of one mutation may depend on the presence of another, a phenomenon known as epistasis [9]. Epistasis is positive when the combined effect of two mutations is greater than expected from their individual effects, and negative when it is smaller [10–12]. Understanding these interactions can help explain adaptive outcomes and may contribute to combinatorial strain engineering. ALE has been used to investigate microbial adaptation for decades [3, 13–16], and mutation data from many ALE experiments have been aggregated in ALEdb (aledb.org) [17], offering an opportunity to investigate epistasis systematically. However, experimental evaluation of every possible mutation pair is impractical, because establishing epistasis requires testing both single mutants and their combination relative to a common genetic background [9, 18, 19]. Therefore, computational prioritization is needed to narrow the search space and select candidate pairs for experimental validation.

Recurrent mutation patterns across ALE endpoints (final samples from lineages evolved in parallel) provide a basis for prioritizing gene pairs for experimental investigation of epistasis [1, 17, 20, 21]. Two patterns of particular interest are co-occurrence, in which two genes are mutated together more often than expected by chance, and mutual exclusivity, in which they are mutated together less often than expected [22]. These patterns have been widely studied in cancer genomics, where co-occurrence may suggest compatibility or synergy, whereas mutual exclusivity may reflect alternative pathways, functional redundancy, or genetic incompatibility [23– 25]. Previous ALE studies have applied descriptive association measures such as Pearson correlation and mutual information to identify and rank such patterns [6, 26, 27]. Although both measures can be used as test statistics, their scores alone do not indicate how unlikely an observed association would be under independence. Statistical association testing addresses this question by comparing the observed score or overlap with an appropriate null distribution, with multiple-testing correction applied when many pairs are screened. These tests have also been applied to ALE data. For example, Fisher’s exact test was used to identify mutual exclusivity between mutations in *rho* and the *rpoBC* operon during high-temperature adaptation of *Escherichia coli B* [20], while a simulation-based mutual-information test identified candidate genetic interactions in yeast, including one pair that was subsequently supported experimentally [21]. These studies illustrate how association testing can identify candidate genetic interactions in ALE mutation data, providing a basis for systematic screening of both co-occurrence and mutual exclusivity across a broad ALE compendium. Machine-learning methods have also been developed to predict epistasis in protein engineering using deep mutational scanning data, evolutionary sequence information, and structural or biophysical priors [28, 29]. For ALE-derived mutation pairs, however, the limited availability of experimentally measured epistatic effects as training labels constrains supervised model training. In this setting, statistical association testing provides a practical approach to prioritizing gene pairs for experimental investigation without requiring such labels.

The results of these statistical association tests depend on the assumptions of null models. A commonly used method for co-occurrence and mutual exclusivity analysis is Fisher’s exact test. When evaluating a gene pair, Fisher’s exact test does not account for differences in the total number of mutated genes among endpoints. Some endpoints may carry mutations in many genes, whereas others carry mutations in only a few, even when they are derived from the same clonal ancestor under the same experimental conditions. Stochastic mutation occurrence, competition among evolving lineages, and hitchhiking of linked mutations can contribute to this variation [30]. Such variation is observed within individual experiments in our ALE compendium (**Fig. 1F**). These differences can affect how often two genes are expected to be mutated together by chance. In cancer genomics, simulations have shown that ignoring differences in the overall number of genomic alterations among tumor samples can produce false-positive co-occurrences and reduce sensitivity for detecting mutual exclusivity [22]. DISCOVER, a statistical method developed for co-occurrence and mutual exclusivity analysis in cancer genomics, addresses such sample heterogeneity by allowing the alteration probability of a gene to vary among individual samples. Applied to ALE data, it estimates a mutation probability for each gene in each endpoint, accounting for both how frequently each gene is mutated and the total number of mutated genes in each endpoint. Pairwise overlap is then evaluated under a Poisson-binomial null model [22].

**Figure 1.**
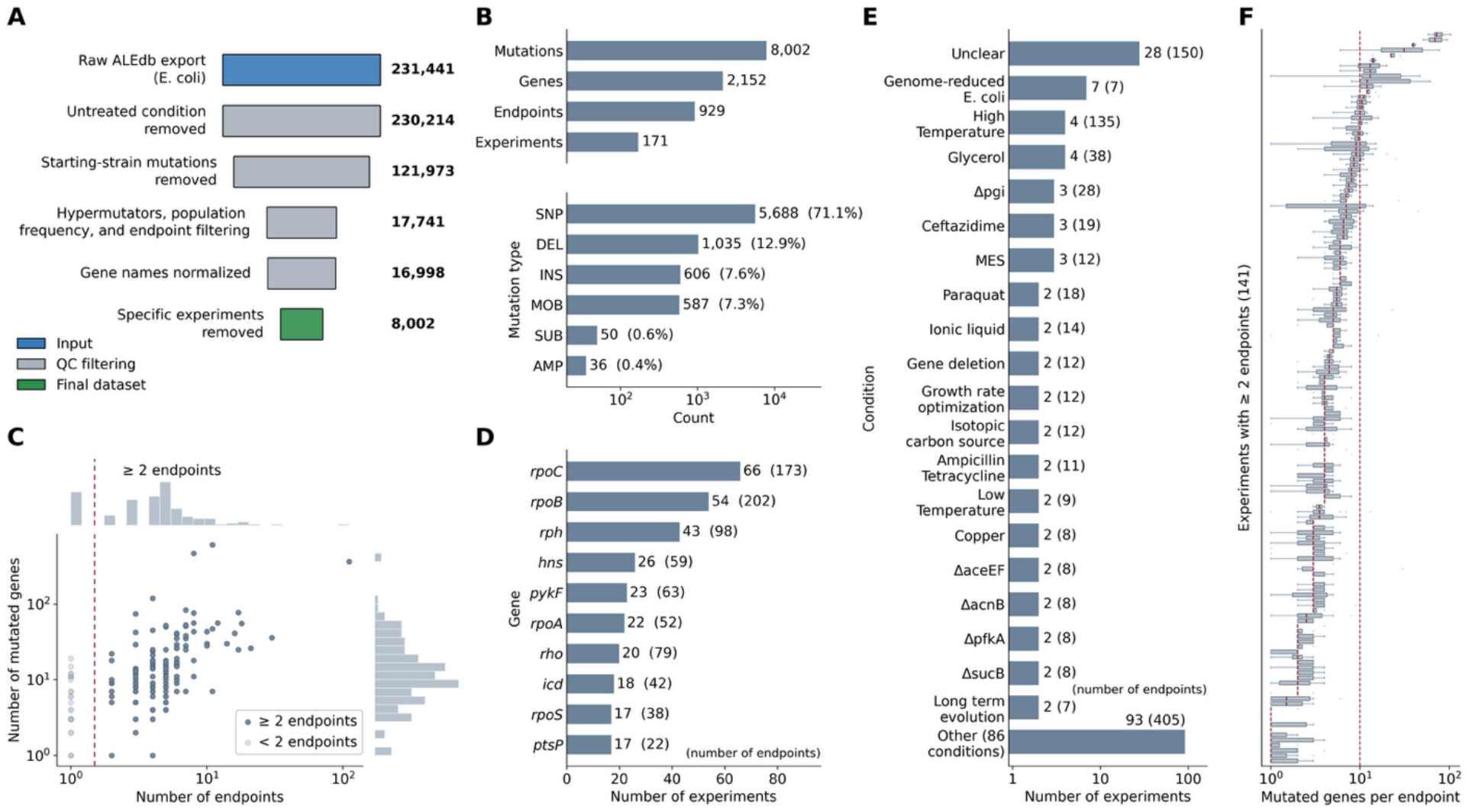
Overview of the curated *Escherichia coli* ALE mutation dataset. **(A)** Sequential quality-control filtering. Bar widths are proportional to the logarithm of the number of mutation records retained at each step, with exact counts shown at right. Colors indicate the input dataset (blue), intermediate filtering steps (gray), and final dataset (green). Filtering criteria are described in **Methods. (B)** Composition of the final dataset. Upper: total numbers of mutations, mutated genes, endpoints, and experiments. Lower: mutation counts by type, with percentages of all mutation records in parentheses. Mutation types are SNP (single-nucleotide polymorphism), DEL (deletion), INS (insertion), MOB (mobile-element insertion), SUB (multi-base substitution), and AMP (amplification). **(C)** Number of mutated genes versus number of endpoints for all 171 experiments after QC, with one point per experiment and marginal histograms. Only genes mutated in at least two endpoints were tested, because a gene mutated in a single endpoint shows no recurrence. Experiments with one endpoint therefore were excluded. The dashed line separates them: dark blue points represent the 141 experiments with at least two endpoints, and light blue points represent the remaining 30. **(D)** The ten most recurrently mutated genes, ranked by the number of experiments in which they are mutated. Values in parentheses indicate the number of endpoints in which each gene is mutated. Ties are resolved by endpoint count. **(E)** The 20 condition categories represented by the most experiments. Values in parentheses indicate endpoint counts for each category. The remaining 86 categories are grouped as ‘Other.’ **(F)** Distribution of mutated genes per endpoint within each of the 141 experiments containing at least two endpoints. Experiments are ordered by their median. The dashed line marks 10 mutated genes per endpoint as a visual reference. Boxes show medians and interquartile ranges (IQRs). Whiskers follow the 1.5×IQR rule, with values beyond the whiskers plotted individually.

We applied DISCOVER alongside Fisher’s exact test as complementary screens to assess the sensitivity of gene-pair associations to null-model assumptions.

In this study, we developed a framework to identify candidate gene pairs for experimental investigation of epistasis and applied it to a curated compendium of 141 *E. coli* ALE experiments from ALEdb. The framework applies Fisher’s exact test and DISCOVER as complementary screens for co-occurrence and mutual exclusivity separately within each experiment, using the same binary gene-by-endpoint matrices. Because ALE endpoints frequently carry different mutations within the same gene, we aggregate mutations at the gene level, categorizing a gene as mutated in an endpoint, and disregarding the specific sequence change. This captures recurrence across distinct mutations within the same gene at the cost of sequence-change resolution. The workflow recovers previously reported mutation patterns, including the mutual exclusivity between mutations in *rho* and RNA polymerase genes during high-temperature adaptation [20], providing a biologically characterized reference for interpreting the statistical results. We examine how null-model assumptions affect statistical support for candidate associations and interpret the results alongside endpoint metadata and mutation-level evidence. Significance under both tests indicates robustness to the choice between these two null models. We also perform a power analysis for Fisher’s exact test to estimate the number of independent endpoints needed to achieve 80% power for detecting gene-pair co-occurrence and mutual exclusivity under pre-specified scenarios. The results suggest that well-powered screening may require more independent endpoints than most experiments in the compendium contained. To automate gene-pair screening across experiments and support reproducible execution and reuse across species and datasets, we implement the framework as a configurable Nextflow pipeline with containerized processes. We release the pipeline and report candidate gene pairs and power estimates to support experimental investigation of epistasis and future ALE study design.

## Results

### A curated compendium of *E. coli* ALE mutations from ALEdb

Aggregated mutational data were retrieved from ALEdb and subjected to sequential quality-control (QC) filtering (**Fig. 1A; Methods**). Restricting the export to *E. coli* leaves 231,441 mutation records. After QC, the resulting compendium comprises 8,002 mutations affecting 2,152 genes across 929 endpoints from 171 experiments (**Fig. 1B**). Each endpoint represented the final sampled state of an ALE (an independently evolved lineage). Single-nucleotide polymorphisms accounted for 71.1% of mutations, followed by deletions (12.9%), small insertions (7.6%), and mobile-element insertions (7.3%), with multi-base substitutions (0.6%) and amplifications (0.4%) making up the remainder.

The number of endpoints and the number of mutated genes varied substantially among experiments (**Fig. 1C**). Of the 171 experiments, 141 contained at least 2 endpoints. The remaining 30 contained a single endpoint and were excluded from pairwise analysis, because candidate genes were required to be mutated in at least 2 endpoints within an experiment (**Methods**). Most experiments were concentrated at the lower end of both distributions, with a median of 4 endpoints per experiment across all 171.

Certain genes were recurrent targets of mutation across experiments (**Fig. 1D**). The RNA polymerase genes *rpoC* and *rpoB* were mutated in 66 and 54 experiments, respectively, corresponding to 173 and 202 endpoints. The most recurrent targets included genes involved in transcriptional machinery and regulation, such as the RNA polymerase genes *rpoA, rpoB*, and *rpoC*, the transcription termination factor gene *rho*, and *hns*, which encodes the global regulator H-NS. RNA polymerase genes are recurring targets in *E. coli* laboratory evolution across diverse conditions [31, 32]. However, widespread recurrence was restricted to a minority of genes. Of the 2,152 mutated genes, 1,301 (60%) were mutated in a single experiment and only 143 (7%) in 5 or more.

The compendium covered diverse experimental conditions, including growth on different carbon sources, exposure to chemical stresses, and high-temperature adaptation (**Fig. 1E**). Examples included experiments involving glycerol, xylose, paraquat, and antibiotics. However, 28 experiments comprising 150 endpoints were assigned to the “Unclear” category because their selective pressure could not be resolved from the available metadata (**Methods**). These unresolved annotations limit condition-specific interpretation of mutation patterns and gene-pair associations.

Endpoints differed considerably in the number of mutated genes they contained (**Fig. 1F**). Across the 899 endpoints from the 141 experiments, the median was 5 mutated genes per endpoint (mean, 7.7), and 80% contained fewer than 10. The distribution was strongly right-skewed, reaching a maximum of 103 mutated genes in a single endpoint. Between experiments, the median number of mutated genes per endpoint ranged from 1 to 72. Within individual experiments, the number of mutated genes also varied among endpoints. In 60 of the 141 experiments, the endpoint with the most mutated genes contained at least 2.5 times as many mutated genes as the endpoint with the fewest. In 28 of these experiments, this ratio was at least 5. This within-experiment variation motivated applying DISCOVER, which accounts for differences among endpoints in the number of mutated genes.

### Fisher’s exact test and DISCOVER differ in null-model structure and dispersion

Both tests were applied to the same data (**Fig. 2A**). After quality control, a binary gene-by-endpoint matrix was constructed for each experiment. Candidate pairs were formed from genes each mutated in at least two endpoints within that experiment. To reduce the number of tests or improve statistical power, previous studies have limited testing to genes or operational units with at least a specified number of mutations [20–22]. However, a single fixed threshold does not account for the variation in experiment size across our compendium (2-111 endpoints). We therefore applied a testability filter separately for each method (Fisher’s exact test and DISCOVER) and direction (co-occurrence and mutual exclusivity). Within each experiment, the testability filter excluded pairs that could not yield a raw *p*-value below 0.05, even at the most extreme overlap allowed by the corresponding null model. The one-sided *p*-value at this extreme is the minimum attainable *p*-value, *p*_*min*_(**Methods**). Pairs passing each filter were tested using the corresponding method and direction, and *q*-values were obtained by Benjamini-Hochberg correction applied separately within each experiment, method, and direction.

**Figure 2.**
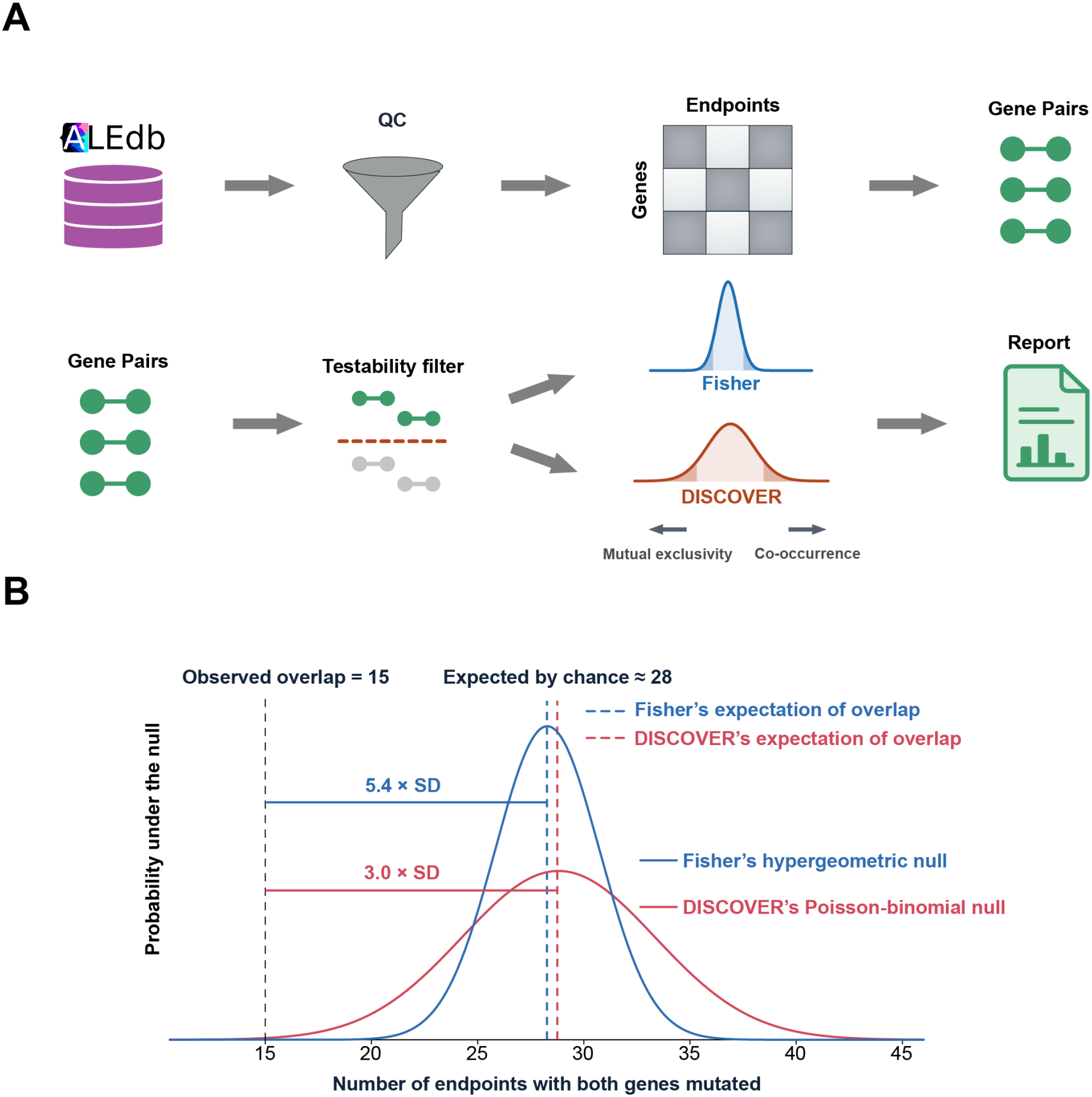
Analysis pipeline and comparison of the Fisher and DISCOVER null distributions for one gene pair. **(A)** Analysis pipeline. After quality control, mutation data from ALEdb are assembled into a binary genes-by-endpoints matrix for each experiment. Candidate pairs comprise two genes each mutated in at least two endpoints within that experiment. Testability is then assessed separately for each method and direction (co-occurrence and mutual exclusivity) using the minimum attainable one-sided *p*-value under the corresponding null distribution. Pairs with *p*_*min*_≥ 0.05 are excluded from further significance testing in that direction (**Methods**). Testable pairs are evaluated under the corresponding method and direction. Benjamini-Hochberg correction is applied within each experiment, method, and direction over the corresponding testable pairs to obtain *q*-values. **(B)** Null distributions of overlap (the number of endpoints in which both genes are mutated) for the *rho-rpoB* pair from the 42.2 °C *E. coli* ALE experiment. Across 111 endpoints, *rho* and *rpoB* are mutated in 43 and 73 endpoints, respectively, with both genes mutated in 15. Fisher’s exact test fixes the observed number of endpoints in which each gene is mutated, giving a hypergeometric null with mean 28.3. The observed overlap lies 5.4 standard deviations below expectation (*p* = 6 × 10^−8^, *q* = 1.5 × 10^−5^). DISCOVER allows these numbers to vary according to estimated gene- and endpoint-specific mutation probabilities. The resulting Poisson-binomial null has mean 28.8 and a larger variance, under which the same observed overlap lies 3.0 standard deviations below expectation (*p* = 1 × 10^−3^, *q* = 0.054). The pair is therefore significant only under Fisher’s exact test at a 5% false discovery rate threshold. Curves are normal approximations shown for visualization.

**Figure S1** illustrates testability using the *glmS-rho* pair in the 42C_Tenaillon experiment. Across 111 endpoints, *rho* was mutated in 43 and *glmS* in 3, with no overlap. Zero overlap is the smallest possible overlap under either null model, but the corresponding one-sided *p-*values were 0.23 under Fisher’s exact test and 0.29 under DISCOVER. These minimum attainable *p*-values exceeded the raw *p*-value cutoff of 0.05. The pair was therefore classified as untestable for mutual exclusivity under both methods and excluded from testing in that direction (mutual exclusivity). It was also untestable for co-occurrence under Fisher’s exact test, but testable under DISCOVER for co-occurrence. Although the pair was testable for co-occurrence under DISCOVER, its observed 0 overlap gave a raw *p*-value of 1. Thus, passing the testability filter establishes that significance is attainable, but does not imply that the observed association is significant.

The two methods differ in two aspects. First, Fisher’s exact test does not account for heterogeneity among endpoints in the total number of mutated genes. DISCOVER accounts for this heterogeneity by estimating a mutation probability for each gene in each endpoint, with the expected number of mutated genes per endpoint constrained to match the observed count. Second, Fisher’s exact test conditions on the observed numbers of endpoints in which each gene is mutated, holding these numbers fixed. DISCOVER instead allows these numbers to vary under the null according to its estimated mutation probabilities. The overlap count follows a hypergeometric distribution under Fisher’s exact test and a Poisson-binomial distribution under DISCOVER [22, 33].

To help determine which aspect of these null models contributed to their different results, we additionally evaluated a binomial model as a diagnostic step. The binomial model assumes that each gene has a constant mutation probability across endpoints and, like Fisher’s exact test, does not account for heterogeneity among endpoints in the total number of mutated genes. Unlike Fisher’s exact test, however, it does not condition on the observed numbers of endpoints in which the two genes are mutated, allowing these counts to vary under the null as in DISCOVER. The binomial and Fisher nulls have the same expected overlap. The binomial model therefore provides an intermediate comparison in which Fisher’s fixed-margin conditioning is removed without introducing DISCOVER’s endpoint-specific mutation probabilities.

These differences in null-model structure also affect the dispersion of the overlap distribution. Fisher’s null holds the number of mutated endpoints for each gene fixed at its observed value, whereas DISCOVER allows these counts to vary according to the estimated gene- and endpoint-specific mutation probabilities. Across 10,290 distinct experiment-specific gene pairs testable by both methods, DISCOVER’s null variance exceeded Fisher’s for every pair, with a variance ratio ranging from 1.12 to 9.92 and a median of 1.36.

The *rho-rpoB* pair from the 42.2 °C *E. coli* ALE experiment [20] provides a concrete example. Across 111 endpoints, *rho* and *rpoB* were mutated in 43 and 73 endpoints, respectively, with both genes mutated in 15 endpoints. The corresponding null distributions are shown in **Fig. 2B**. They had similar expected overlaps, 28.3 for Fisher’s exact test and 28.8 for DISCOVER, but differed substantially in variance. Under Fisher’s exact test, the observed overlap (15) is 5.4 standard deviations below expectation, giving *p* = 6 × 10^−8^, and after multiple-testing correction *q* = 1.5 × 10^−5^. Under DISCOVER, whose null standard deviation was 1.9 times larger (3.5 times in variance), the same overlap was 3.0 standard deviations below expectation, giving *p* = 1 × 10^−3^ and *q* = 0.054. Thus, the pair was significant under Fisher’s exact test but not under DISCOVER at a 5% false discovery rate threshold.

### Fisher’s exact test and DISCOVER identify different sets of significant gene pairs

The two methods yielded different sets of testable gene pairs. Overall, 15,398 experiment-specific gene pairs across 94 experiments were testable by at least one method in at least one direction (co-occurrence or mutual exclusivity). For co-occurrence, 10,171 pairs from 29 experiments were testable by Fisher’s exact test, compared with 15,216 pairs from 83 experiments by DISCOVER. For mutual exclusivity, 770 pairs from 25 experiments were testable by Fisher’s exact test, compared with 969 pairs from 64 experiments by DISCOVER.

At a 5% false discovery rate threshold, Fisher’s exact test identified 74 significant gene pairs across 12 experiments, comprising 36 co-occurrences and 38 mutual exclusivities. DISCOVER identified 5 significant pairs, all for mutual exclusivity, across 4 experiments. All 5 pairs were also significant by Fisher’s exact test. Of the 141 experiments containing at least 2 endpoints, 129 yielded no significant pair under either method (**Fig. 3A**). Among the remaining 12 experiments, 3 yielded only significant co-occurrences, 5 yielded only significant mutual exclusivities, and 4 yielded both.

**Figure 3.**
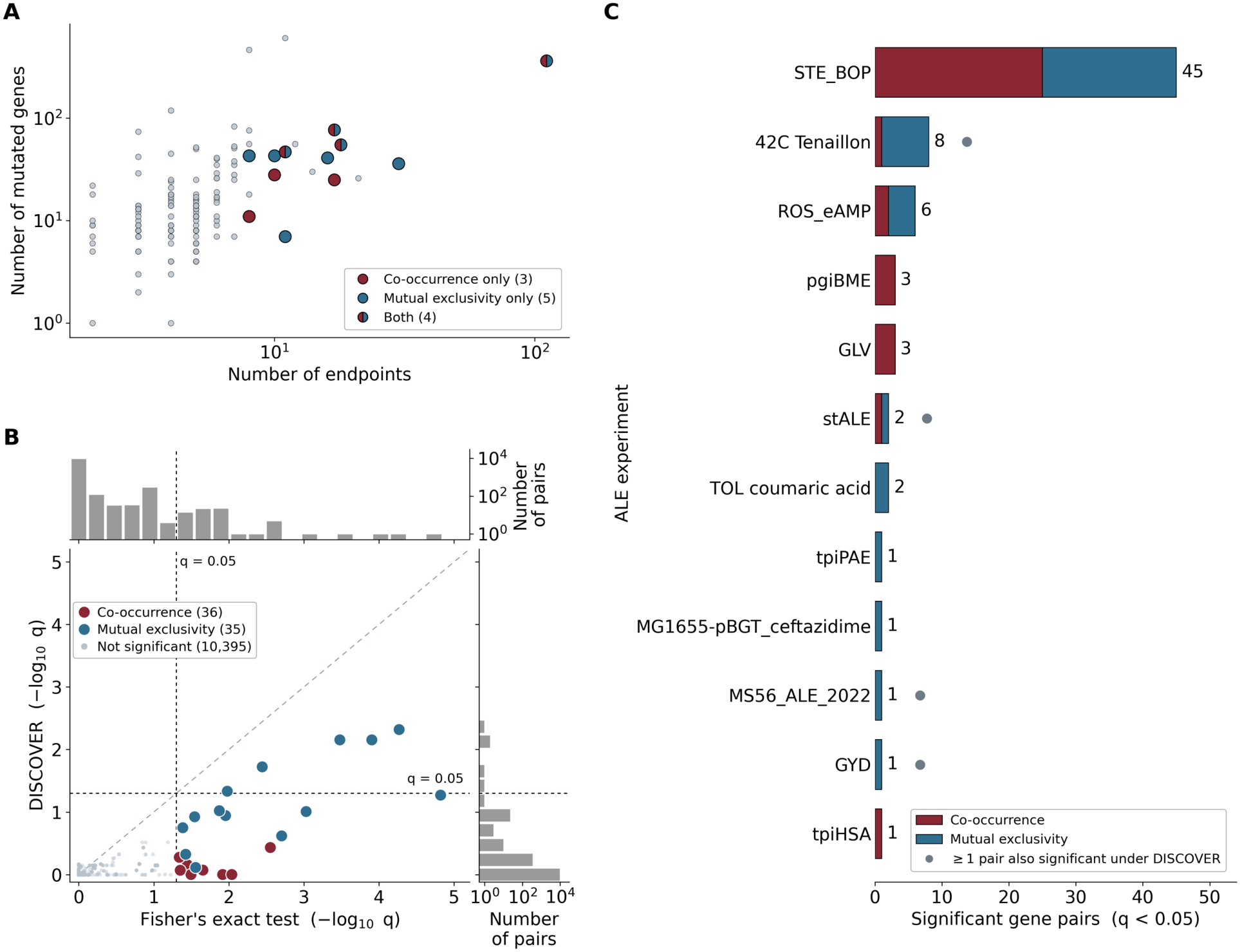
Pairwise co-occurrence and mutual exclusivity analysis results across the *E. coli* ALE compendium. Gene pairs were tested separately within each experiment using one-sided Fisher and DISCOVER tests. Benjamini-Hochberg correction was applied within each experiment, method, and direction over testable pairs, with *q* < 0.05 defining significance. **(A)** Number of mutated genes versus number of endpoints for the 141 experiments with at least two endpoints, with one point per experiment. Gray points represent 129 experiments with no significant pair. Colored points represent the 12 experiments with at least one: red for co-occurrence only, blue for mutual exclusivity only, and split markers for both directions. **(B)** Comparison of *q*-values for pairs testable by both methods in the same direction. Testability required a minimum attainable one-sided *p*-value below 0.05 (**Methods; Fig. S1**). Each experiment-specific pair appears once per shared testable direction, yielding 10,171 co-occurrence and 295 mutual exclusivity pairs. Pairs testable in both directions appear twice. Axes show ™log_10_(*q*). Gray points represent 10,395 pairs that were nonsignificant under both methods. Red and blue points represent the 36 co-occurrence and 35 mutual exclusivity pairs significant under at least one method. Points with identical *q*-values overlap, including 8,836 comparisons at *q* = 1 under both methods. Marginal histograms show the *q*-value distributions, with counts on logarithmic scales. Three Fisher-significant pairs are omitted because they were untestable for mutual exclusivity under DISCOVER. Dashed lines mark *q* = 0.05, and the diagonal indicates equal *q*-values. All 71 plotted Fisher-significant comparisons lie below the diagonal, indicating larger DISCOVER *q*-values. **(C)** Numbers of Fisher-significant gene pairs in the 12 experiments with significant associations, with bars stacked by direction. Gray dots identify the four experiments containing at least one DISCOVER-significant pair.

DISCOVER returned a larger *q*-value than Fisher’s exact test for 71 Fisher-significant pairs (**Fig. 3B**). The remaining 3 pairs were untestable for mutual exclusivity under DISCOVER and were therefore omitted from the figure. The two methods produced similar expected overlaps, with a median absolute difference of 0.05 endpoints across these 71 pairs. In contrast, as discussed above, DISCOVER’s null variance exceeded Fisher’s for all pairs testable by both methods in the same direction. This greater null dispersion was associated with the larger DISCOVER raw *p*-values among the Fisher-significant pairs.

We used the binomial test introduced above as a diagnostic comparison to assess whether accounting for differences among endpoints in the total number of mutated genes was the main source of the Fisher-DISCOVER difference. Its testable sets were similar to those of DISCOVER (15,235 co-occurrence and 837 mutual exclusivity pairs, versus 15,216 and 969 for DISCOVER) and at a 5% false discovery rate threshold, the binomial test identified no significant co-occurrences and 3 significant mutual exclusivity pairs. All were also significant under DISCOVER. Among the 74 Fisher-significant pairs, the median

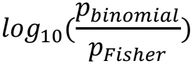

was 1.261, whereas the median

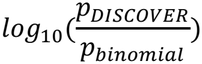

was -0.048. For every one of these 74 pairs, the binomial raw *p*-value was closer to DISCOVER’s than to Fisher’s on the log_10_(*p*) scale. Because the binomial and Fisher nulls share the same expected overlap, while the binomial and DISCOVER nulls both allow the numbers of endpoints in which each gene is mutated to vary, these results support differences in null dispersion as the main explanation for the Fisher-DISCOVER *p*-value discrepancy among these 74 pairs, rather than DISCOVER’s modeling of endpoint heterogeneity in the total number of mutated genes. The remaining differences between the binomial test and DISCOVER show that endpoint-specific mutation probabilities still affected testability and which pairs reached significance after multiple-testing correction.

The 74 significant pairs identified by Fisher’s exact test were unevenly distributed across experiments. The STE_BOP experiment contributed 45, accounting for 61% of all significant pairs (**Fig. 3C**). This experiment included 26 candidate genes across 11 endpoints and yielded 45 significant pairs: 25 for co-occurrence and 20 for mutual exclusivity. The 42C_Tenaillon experiment, which contained the most endpoints in the compendium, contributed 8 significant pairs.

Significant pairs were sparse under both methods despite the large testable sets, although the number of significant pairs alone does not establish low statistical power. Testability and power address different questions. Testability determines whether a pair can attain the specified raw *p*-value threshold under a given null distribution, whereas power is the probability of rejecting the null when an association of a specified magnitude is present. To estimate the number of independent endpoints needed to achieve 80% power for detecting gene-pair co-occurrence and mutual exclusivity, we performed a power analysis for Fisher’s exact test. In the power analysis, each gene was assigned a mutation probability of 0.15 per endpoint, and endpoints were independent. This value closely matched the empirical marginal proportions of mutated endpoints among genes in Fisher-testable pairs, for which the pair-weighted mean was 0.149. For a single pair tested at *p* < 0.05 without multiple-testing correction, 70 endpoints were required to achieve 80% power for co-occurrence at odds ratio (OR) = 10, compared with 215 endpoints required for mutual exclusivity at OR = 0.1 (**Fig. S2A, B**). Under this model, only 42C_Tenaillon, with 111 endpoints, exceeded the 70-endpoint benchmark, and no experiment reached the 215-endpoint benchmark.

To assess how multiple-testing correction affects power when screening many gene pairs, we used Bonferroni thresholds based on the numbers of Fisher-testable pairs in each direction in 42C_Tenaillon:

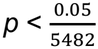

for co-occurrence and

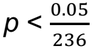

for mutual exclusivity. Under these thresholds, 80% power required 231 endpoints for co-occurrence and 590 endpoints for mutual exclusivity (**Fig. S2A, B**). None of the 141 analyzed experiments contained enough endpoints to reach either of these 80% power benchmarks under the reference scenarios. Across a broader range of pre-specified odds ratios, the endpoints required to achieve a given power increased sharply as the odds ratios approached 1 (**Fig. S2C**).

### Four experiments contain gene pairs significant under both Fisher’s exact test and DISCOVER

Across 4 experiments, 5 gene pairs were significant for mutual exclusivity under both Fisher’s exact test and DISCOVER (**Fig. 4A**). This panel also shows the 7 additional pairs significant only under Fisher’s exact test in these experiments, giving 12 pairs in total. Panels **B-E** show endpoint-level mutation patterns for the genes involved in these pairs, with one panel for each experiment. The 12 pairs are discussed below as 3 biological cases: high-temperature adaptation (42C_Tenaillon), glycerol adaptation (GYD), and mutual exclusivity between RNA polymerase subunit genes.

**Figure 4.**
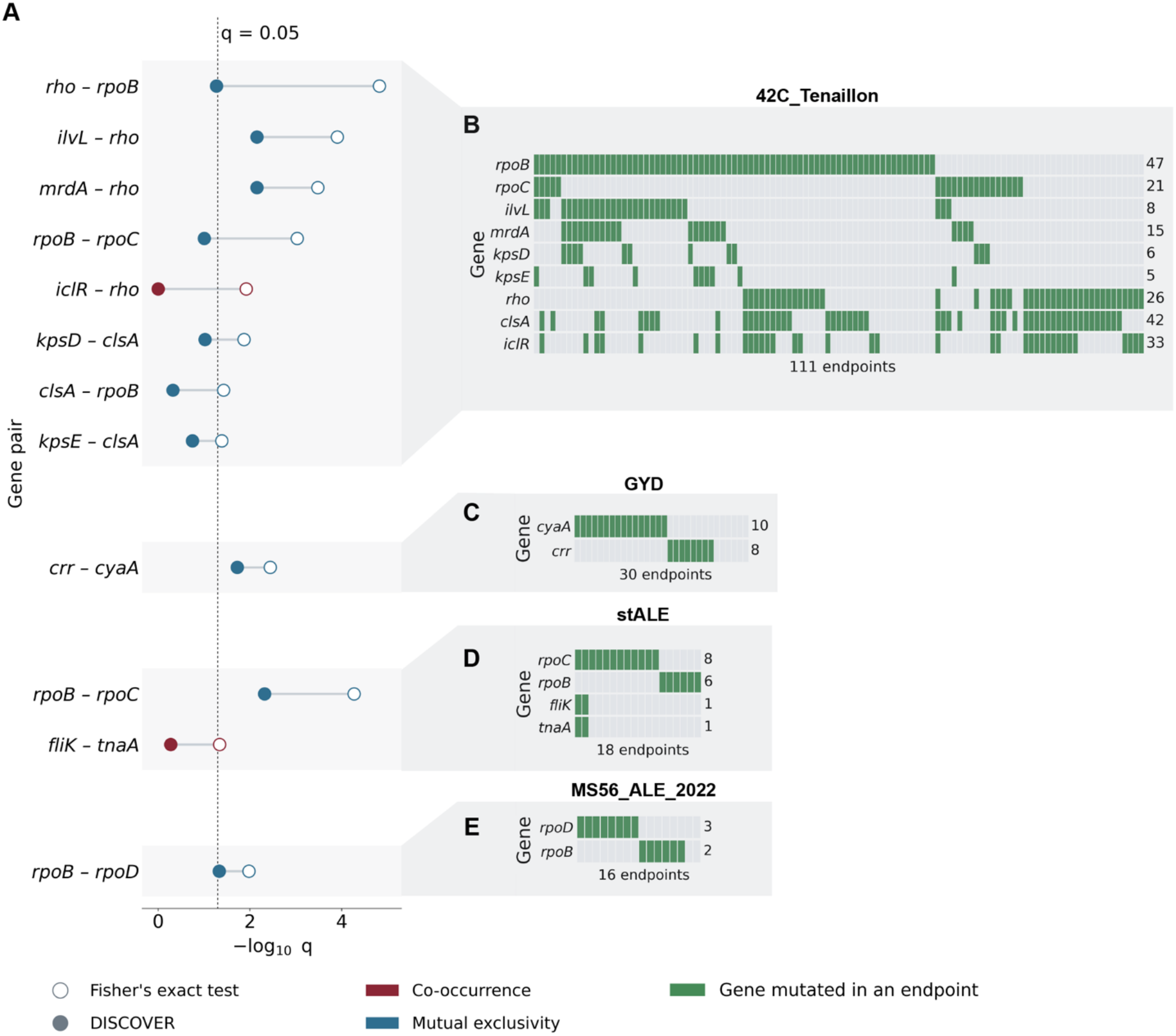
DISCOVER identifies a subset of Fisher-significant gene pairs, with their corresponding endpoint-level mutation patterns. Four experiments contain at least one gene pair significant under both Fisher’s exact test and DISCOVER. Experiments are ordered by number of endpoints. (**A**) The 12 pairs significant by Fisher’s exact test in these 4 experiments are shown, grouped by experiment. Five of them are also significant by DISCOVER. Open circles denote Fisher’s exact test, while DISCOVER circles are filled. Red indicates co-occurrence and blue indicates mutual exclusivity. q-values are Benjamini-Hochberg corrected within each experiment, method, and direction over all pairs testable in that direction. The dashed line marks *q* = 0.05, with values to the right indicating significance. All 5 pairs significant under both methods show mutual exclusivity. (**B–E**) Endpoint-level mutation patterns for the same four experiments. Rows show the genes belonging to the pairs displayed in panel A for each experiment. Columns represent all endpoints in the experiment, ordered by mutation pattern. Green indicates that a gene is mutated in an endpoint. The number at the right of each row indicates the number of distinct mutations recorded for that gene in the experiment. A value of 1 therefore indicates that all mutated endpoints carry the same mutation. Endpoints with no mutations in any displayed gene appear as empty columns at the right: 6 of 30 in GYD, and 2 of 16 in MS56_ALE_2022. In the 42C_Tenaillon experiment and stALE, every endpoint contains a mutation in at least one displayed gene.

### The high temperature tolerance evolution experiment (42C_Tenaillon)

The 42C_Tenaillon experiment derives from the study by Tenaillon et al., who evolved 115 lines of *E. coli B* at 42.2 °C and sequenced one endpoint clone from each, identifying two competing adaptive routes: one involving the *rpoBC* operon and the other involving the transcription termination factor *rho* [20]. After quality control, our reconstruction of their data retains 111 of the 114 non-hypermutator lines and nearly all mutations in the genes central to these routes: 84 of 87 in *rpoB*, 21 of 21 in *rpoC*, and 43 of 45 in *rho*. Of the 111 retained endpoints, 68 carried mutations in *rpoB* and/or *rpoC* but not *rho*, 22 in *rho* but neither *rpoB* nor *rpoC*, and 21 in both. All 8 Fisher-significant pairs in this experiment involve genes assigned by the source study to one route or the other (**Fig. 4A, B**). Among these 8 Fisher-significant pairs, *ilvL-rho* and *mrdA-rho* are also significant under DISCOVER. The *rho-rpoB* pair is significant only under Fisher’s exact test and falls slightly above the DISCOVER threshold (*q* = 0.054).

The *rho-rpoB* pair showed significant mutual exclusivity under Fisher’s exact test (*q* = 1.5 × 10^−5^), although the exclusivity was not absolute at the gene level. 15 of the 111 lines carry mutations in both genes, but most of these co-mutated endpoints involve non-recurrent *rho* mutations. By contrast, the *rho* mutations I15N and I15F occurred in 16 endpoints, none of which carried a mutation in any RNA polymerase subunit genes. The pattern is symmetric: the *rpoB* mutations, *I966S* and *I966N*, account for 18 lines, and none carries a mutation in *rho*. Repeated attempts to construct a double mutant carrying *rho I15N* and *rpoB I966S*, the most frequent *rho* and *rpoB* mutations in the experiment, were unsuccessful, and all 8 *rho-rpoB* combinations that could be constructed showed negative epistasis at 42.2 °C, with 4 reaching statistical significance [34].

Apart from the *rho-rpoB and rpoB-rpoC pairs*, the remaining 6 Fisher-significant pairs identified in this experiment may reflect the same division between adaptive routes. The *rpoB*-*rpoC* pair is discussed below with other associations between RNA polymerase subunit genes. Mutations in *clsA* and *iclR* are concentrated among endpoints carrying *rho* mutations without a mutation in RNA polymerase subunit genes, whereas mutations in *ilvL, mrdA, kpsD*, and *kpsE* occur only in endpoints that carry mutations in RNA polymerase subunit genes. When reconstructed individually in the ancestral strain, *rho* I15N and I15F produced no measurable fitness benefit and were inferred to be beneficial only in combination with other mutations, whereas two other recurrent *rho* mutations were beneficial on their own [32]. The *iclR-rho* pair is the only significant co-occurrence in this experiment under Fisher’s exact test (*q* = 0.012), although it is not significant under DISCOVER. *González-González et al*. proposed a possible functional explanation for this association: *iclR* mutations might compensate for the reduced *aceBAK* expression observed in *rho* mutants. The functional relationship between *clsA* and *rho*, however, remained unresolved in that study [32].

DISCOVER does not identify the *rho-rpoB* pair as significant (raw *p* = 1.0 × 10^−3^, *q* = 0.054). As described above, the two methods assign similar expected overlaps, but DISCOVER’s broader null distribution makes the same observed overlap less extreme. Gene-level aggregation may further weaken the association: *rho* I15N and I15F never occur alongside a *rpoB* or *rpoC* mutation, whereas 21 of the 27 endpoints carrying other *rho* mutations also carry a mutation in RNA polymerase genes.

### Mutual exclusivity of *cyaA* and *crr* in the glycerol adaptation experiment (GYD)

In the GYD experiment, *E. coli* evolved with glycerol as the sole carbon source, yielding 30 endpoints. The source study identified *glpK, cyaA*, and *crr* as the 3 most frequently mutated genes and mapped all 3 to the CRP regulon [26]. The authors also reported a negative Pearson correlation between the mutation profiles of cyaA and crr, which they interpreted as evidence of negative epistasis [26]. Although this correlation highlighted the *cyaA-crr* relationship, it did not formally assess how unlikely the observed overlap was under a specified null model. Therefore, we formally tested all testable gene pairs in the GYD experiment for co-occurrence and mutual exclusivity using Fisher’s exact test and DISCOVER. The *crr-cyaA* pair was the only significant pair. Of the 30 endpoints, 16 carried a *cyaA* mutation, and 8 carried a *crr* mutation, but none carried both. The pair was significant for mutual exclusivity under both Fisher’s exact test (*q* = 3.6 × 10^−3^) and DISCOVER (*q* = 0.019), and 24 of the 30 endpoints carried a mutation in one of the two genes (**Fig. 4A, C**).

The products of *crr* and *cyaA* act sequentially in the cAMP-CRP regulatory pathway. *crr* encodes the glucose-specific phosphotransferase system (PTS) protein IIA^Glc^, whereas *cyaA* encodes adenylate cyclase. During growth on glycerol, a non-PTS carbon source, IIA^Glc^ becomes phosphorylated and activates adenylate cyclase, which is involved in the synthesis of cyclic adenosine monophosphate (cAMP). cAMP is a signaling molecule that binds the cAMP receptor protein (CRP), a global transcriptional regulator. The resulting cAMP-CRP complex activates genes involved in glycerol metabolism and other secondary-carbon pathways, thereby relieving carbon catabolite repression (CCR) [35, 36].

Within this regulatory framework, the source study interpreted the *glpK, cyaA*, and *crr* mutations together as a potential route to increasing glycerol metabolism while maintaining repression of other secondary-carbon systems. Specifically, *glpK* mutations may increase glycerol kinase activity by disrupting inhibitory regulation, with many mutations predicted to reduce fructose-1,6-bisphosphate-mediated inhibition, whereas mutations in *cyaA* or *crr* may reduce cAMP synthesis and maintain carbon catabolite repression in glycerol medium [26]. Because cAMP-CRP also contributes to activation of the *glpFKX* operon, reduced cAMP signaling would not be expected to enhance glycerol utilization directly and could also reduce transcription of glycerol-utilization genes [26].

Despite this unresolved effect on glycerol-pathway transcription, mutations in *crr* and *cyaA* are predicted to converge on the same regulatory outcome: reduced cAMP-CRP signaling. Under this model, they represent alternative adaptive routes to a similar regulatory state. Once one route has been taken, mutation of the other gene may provide little additional benefit, offering a plausible explanation for their mutual exclusivity. However, the observed mutual exclusivity does not by itself establish negative epistasis, and experimental validation is still needed.

### Mutual exclusivity between RNA polymerase subunit genes

The remaining 2 of the 5 pairs significant under both methods involve mutual exclusivity between *rpoB* and other RNA polymerase subunit genes. In the stALE experiment, *rpoB* and *rpoC* were mutated in 6 and 12 of 18 endpoints, respectively, but never in the same endpoint (Fisher’s exact test, *q* = 5.4 × 10^-5^; DISCOVER, *q* = 0.005). In the MS56_ALE_2022 experiment, *rpoB* and *rpoD* were mutated in 6 and 8 of 16 endpoints, respectively, and likewise never occurred together (Fisher’s exact test, *q* = 1.0 × 10^-2^; DISCOVER, *q* = 0.047). Mutual exclusivity involving *rpoB* recurred across two independent experiments, although the partner gene differed (**Fig. 4A, D, E**).

RNA polymerase genes are recurrent targets in ALE experiments (**Fig. 1D**), and adaptive mutations in these genes can broadly reprogram transcription [37]. Mutations in different components of the RNA polymerase may therefore produce partially overlapping changes in gene expression. Such overlap could reduce the additional benefit of a second mutation and thereby contribute to the mutual exclusivity patterns.

Mutual exclusivity among genes encoding RNA polymerase components was not consistently observed across experiments, indicating that these patterns may be context dependent. In 42C_Tenaillon, *rpoB* and *rpoD* were mutated together in 19 of 111 endpoints and showed no evidence of mutual exclusivity (*q* = 1.00). The *rpoC-rpoD* pair was likewise nonsignificant. Across the compendium, *rpoB-rpoC* was testable for mutual exclusivity by both methods in 3 experiments.

It was significant under both methods in stALE and under Fisher’s exact test alone in 42C_Tenaillon. The third experiment, with 11 endpoints, was nonsignificant under both methods.

Together, the 5 pairs significant under both methods span 3 biological contexts, but the strength and nature of the supporting evidence differ. In 42C_Tenaillon, *ilvL-rho* and *mrdA-rho* follow the broader division between *rho-* and RNA polymerase-associated adaptive routes. Experimental reconstruction of the *rho-rpoB* pair has demonstrated negative epistasis at 42.2 °C, supporting the biological relevance of this broader division, although it does not directly validate epistasis for *ilvL-rho* or *mrdA-rho* [34]. The mechanistic interpretation of *crr-cyaA* remains experimentally untested, and the *rpoB-rpoC* and *rpoB-rpoD* associations are experiment-specific in the present compendium.

### Most associations significant only under Fisher’s exact test coincide with multiple strain labels or single-mutation recurrence

Of the 74 gene pairs significant under Fisher’s exact test, 5 were also significant under DISCOVER, leaving 69 pairs significant only under Fisher’s exact test. To characterize the difference between these two sets, we examined two features of the deposited metadata and mutation records that are relevant to interpreting mutation patterns. An experiment was classified as having multiple strain labels when its endpoints were recorded under more than one strain name in ALEdb. Single-mutation recurrence was defined as a gene mutated in more than one endpoint, with only one distinct mutation recorded across all affected endpoints.

None of the 5 pairs significant under both methods arose from an experiment with multiple strain labels, and none involved a gene showing single-mutation recurrence. For each of these 5 pairs, both genes were represented by at least 2 distinct mutations across the endpoints in which they were mutated (**Fig. 4B-E**). By contrast, 45 of the 69 pairs significant only under Fisher’s exact test arose from STE_BOP, whose endpoints have multiple strain labels, and 57 of the 69 pairs involved at least one gene showing single-mutation recurrence. 60 pairs showed one or both features (**Fig. 5A**). These two features were present in most pairs significant only under Fisher’s exact test.

**Figure 5.**
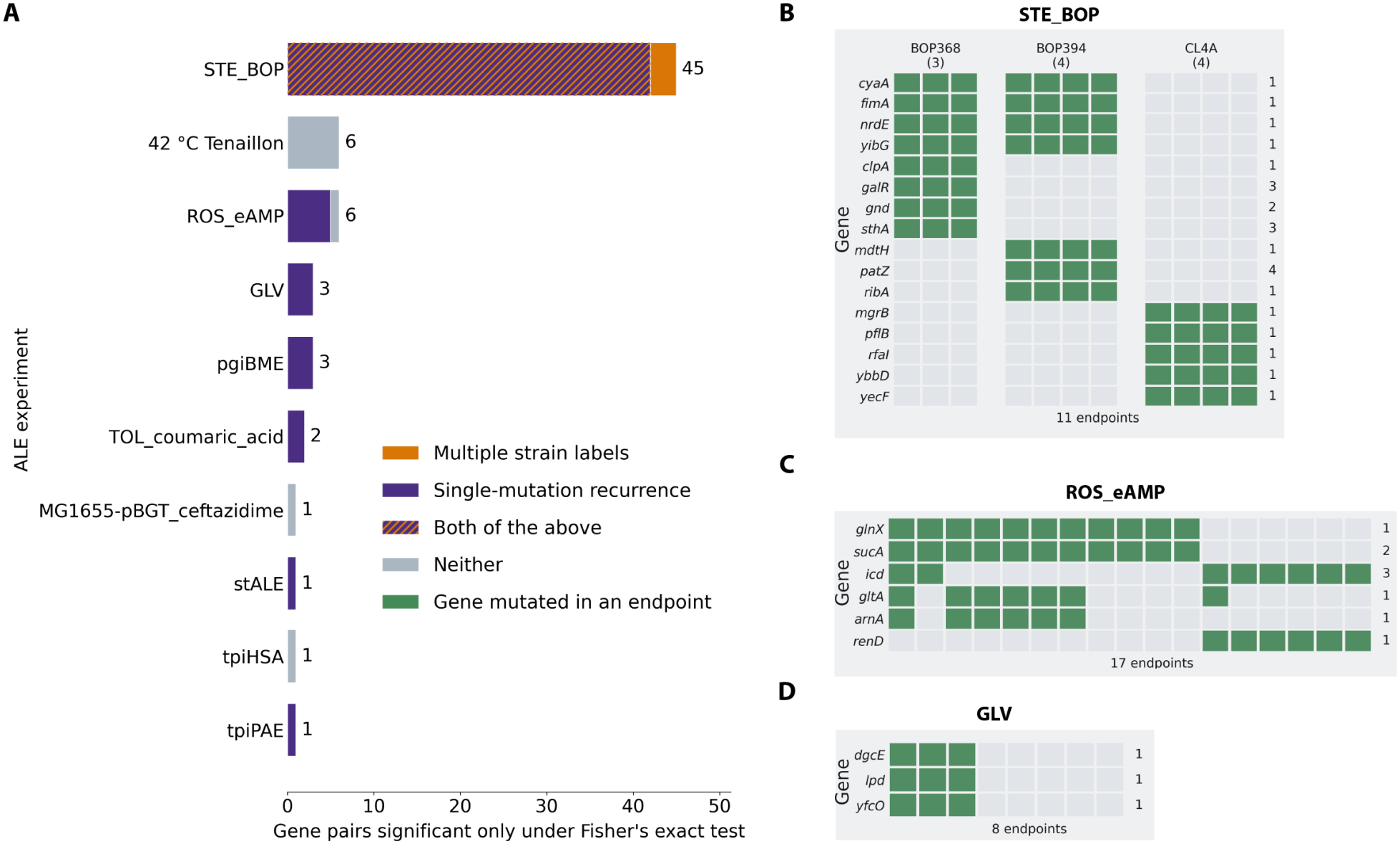
Most Fisher-only significant pairs involve multiple strain labels or single-mutation recurrence. Of the 74 experiment-specific gene pairs significant under Fisher’s exact test, 5 were also significant under DISCOVER. This figure characterizes the remaining 69 pairs. Significance denotes *q* < 0.05 after Benjamini-Hochberg correction within each experiment, method, and direction over testable pairs. (**A**) Number of Fisher-only pairs per experiment, colored by two features of the metadata and mutation records. Amber marks pairs from experiments with more than one endpoint strain annotation. Purple marks pairs involving at least one gene mutated in multiple endpoints, with only one distinct mutation recorded across all affected endpoints (single-mutation recurrence). Purple with amber hatching marks pairs with both features, and gray marks pairs with neither. Of the 69 pairs, 45 come from STE_BOP, whose endpoints have multiple strain labels, 57 involve single-mutation recurrence, 60 show one or both features, and nine show neither. (**B-D**) Mutation patterns for 3 experiments from panel A. Rows represent genes belonging to the pairs in panel A for each experiment, and columns represent all endpoints. Green indicates mutation presence. Numbers at the right give the number of distinct mutations recorded for the gene. A value of one indicates the same mutation in every affected endpoint. In panel **B**, columns are grouped by the strain annotation, with strain labels and endpoint counts above each group. Rows are ordered by their mutation patterns across groups.

In ALEdb, samples are indexed by ALE, flask, isolate, and technical replicate identifiers [17]. The STE_BOP export includes a starting-strain record labeled A0 F0 I1 R1. However, its 11 endpoints carry 3 distinct strain labels: BOP368, BOP394, and CL4A, with 3, 4, and 4 endpoints, respectively. The metadata do not clarify how the starting-strain record relates to these groups or whether all groups share the same ancestral strain.

The 16 genes involved in the 45 Fisher-significant pairs showed 4 group-level mutation patterns: 4 were mutated in all 7 endpoints labeled BOP368 or BOP394, 4 only in BOP368, 3 only in BOP394, and 5 only in CL4A (**Fig. 5B**). All 25 co-occurrence pairs involved genes mutated in identical sets of endpoints, whereas all 20 mutual exclusivity pairs involved genes mutated in nonoverlapping sets. For 12 of the 16 genes, the same mutation was recorded in every affected endpoint (single-mutation recurrence), and 42 of the 45 pairs involved at least one such gene. Analyzing all 11 endpoints together can yield Fisher-significant associations that reflect the endpoint groups, without requiring a genetic interaction between the genes.

Except for pairs from STE_BOP, 15 additional pairs involved single-mutation recurrence. This feature accounted for 5 of the 6 pairs in ROS_eAMP and all pairs in GLV, pgiBME, TOL_coumaric_acid, stALE, and tpiPAE (**Fig. 5A**). In ROS_eAMP, *glnX* is mutated in 11 endpoints, *gltA* in 7, and *arnA* and *renD* in 6. For each of these 4 genes, the same mutation is detected in every affected endpoint (**Fig. 5C**). In GLV, *dgcE, lpd*, and *yfcO* are each mutated in the same 3 endpoints, and the same mutation is detected for each gene in all 3 endpoints (**Fig. 5D**).

Single-mutation recurrence may indicate repeated selection of the same mutation in independent evolutionary lineages [6]. However, this can also arise through inheritance of a mutation that was present before the analyzed lineages diverged, repeated sampling of related lineages, or another source of non-independence among endpoints. The metadata of these experiments do not contain sufficient information to distinguish among these possibilities. We therefore treat single-mutation recurrence as a flag for further examination of mutation origins.

The two features have different implications for interpreting gene-pair associations. Multiple strain labels may indicate genetic-background stratification within an experiment, whereas single-mutation recurrence leaves the independence of mutation origins unresolved. Fisher’s exact test conditions on each gene’s observed mutated-endpoint count and treats endpoints as exchangeable under the null, so stratification or other dependence among endpoints can generate apparent gene-pair associations. DISCOVER estimates gene- and endpoint-specific mutation probabilities and accounts for differences in the total number of mutated genes among endpoints. In the unstratified analysis used here, DISCOVER did not incorporate the recorded strain-label groups or resolve whether repeated occurrences of the same mutation arose independently. Excluding STE_BOP reduced the number of pairs significant under Fisher’s exact test from 74 to 29.

### A configurable pipeline for reproducible pairwise association analysis of ALE data

The previous analyses showed that pairwise results depend on the statistical null model, the testability criterion, and the structure and curation of the input data. To make these choices explicit and reproducible, we implemented the data preparation and pairwise testing as a configurable Nextflow pipeline (**Fig. 6**) [38]. The implementation addresses 4 practical requirements. First, reproducibility: each QC stage, QC report generation, and the pairwise analysis run as separate processes using a Docker image with fixed software versions, providing a consistent software environment across compatible systems. Second, consistency: the same command ran unchanged on a Linux server and on a laptop, producing identical QC tables and significance calls on both systems. Third, efficient reanalysis: the -resume option allows cached results from unchanged tasks to be reused, avoiding unnecessary re-computation when analysis settings are modified. Fourth, traceability: each run records its parameters, the code version, and the container image, and Nextflow can report execution time and memory usage for individual tasks. A complete *E. coli* analysis took approximately 2 minutes on a Linux server (4 virtual CPUs with 32 GB of RAM) and 2.5 minutes on the laptop (8-core CPU with 16 GB of RAM). The pipeline, its configuration files, and the public input data are available at https://github.com/Multiomics-Analytics-Group/ale-cooccurrence-mutual-exclusivity-nf.

**Figure 6.**
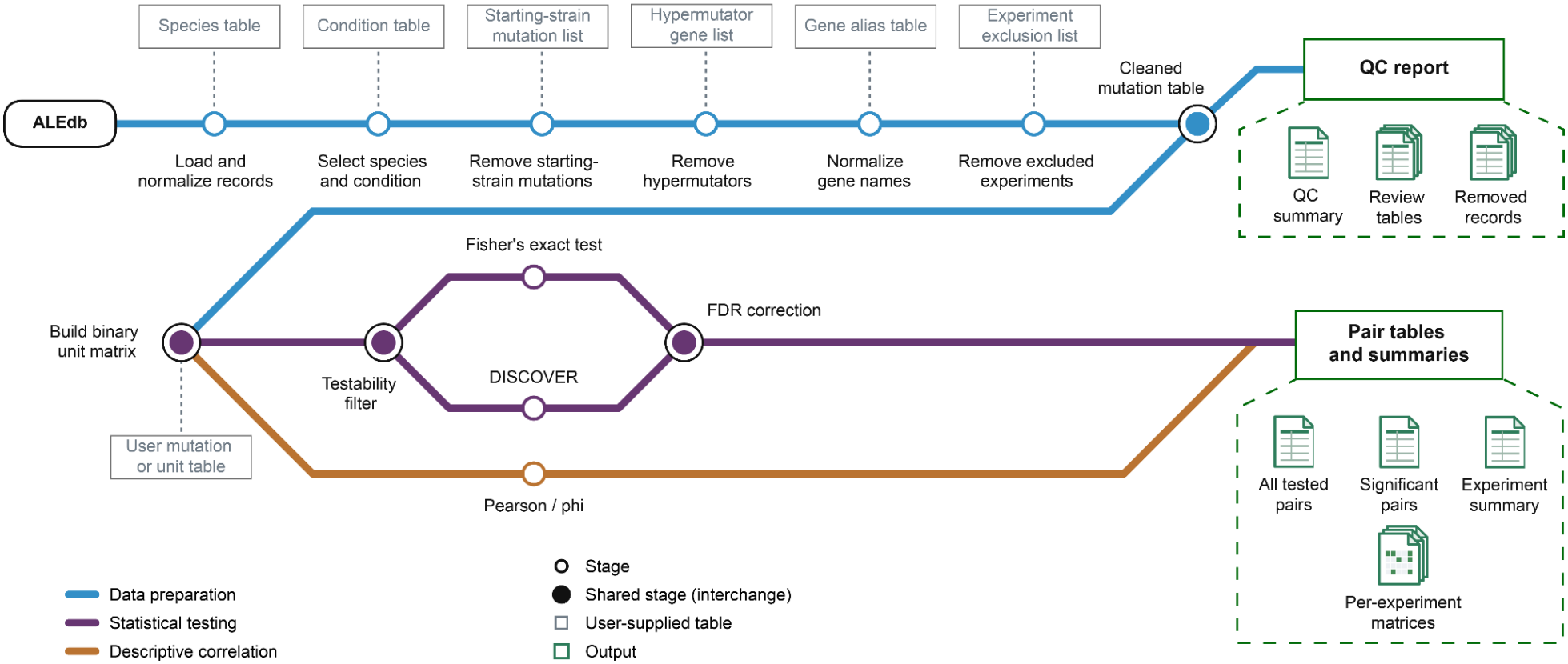
Metro map of the configurable analysis pipeline. Lines show the routes data can take through the workflow, and stations mark processing stages. Double-ringed stations are shared by more than one route. Blue denotes data preparation. Six quality-control stages transform the raw ALEdb export into a cleaned mutation table. The quality-control report records the number of mutations entering and leaving each stage, providing the counts shown in Fig. 1A. Squares connected by dashed lines denote user-supplied tables. These tables provide normalization rules, starting-strain mutations, hypermutator genes, gene aliases, and experiment exclusions, keeping dataset-defining decisions outside the code. The matrix stage can also accept a user-supplied mutation or unit table. This allows data from sources other than ALEdb to enter the workflow and allows genes to be replaced by operons, pathways, or other user-defined test units. Purple denotes statistical testing. One binary test-unit-by-endpoint matrix is built for each experiment, with genes used as the test units in this study. A testability filter is applied separately for each method and direction and removes pairs whose most extreme attainable overlap cannot yield *p* < 0.05 The remaining pairs are tested for co-occurrence and mutual exclusivity by Fisher’s exact test and DISCOVER. Benjamini-Hochberg correction is then applied separately within each experiment, method, and direction over all pairs testable in that direction. Amber denotes descriptive correlation. A Pearson correlation is computed for each pair as an effect size and reported without a significance test, so this route passes through neither the testability filter nor the correction stage. Green rectangles denote outputs.

Data preparation converts the raw ALEdb export into a cleaned mutation table through six ordered stages. These stages load the ALEdb mutation calls and metadata, select the species, and remove the condition groups named in the configuration (for *E. coli*, untreated group). They then remove starting-strain mutations and hypermutator lineages, normalize gene names, and remove excluded experiments. Each stage writes its output, and every filtering stage also records the entries it removes. A quality-control report and a summary file for each stage record the number of mutations entering and leaving each stage, providing the counts shown in **Fig. 1A**. The cleaned output retains the endpoint metadata needed to inspect features such as multiple strain labels, while decisions about exclusion or stratification remain under user control.

From the cleaned table, the pipeline builds one binary matrix of test units by endpoints for each experiment. The test unit is specified by a column name, allowing genes to be replaced by operons, pathways, or other user-defined groupings without code changes. The analysis pipeline (statistical testing part) also accepts a user-supplied mutation table, allowing data from sources other than ALEdb to enter the workflow. For each method and direction, pairs that cannot pass the testability filter were excluded before testing. The remaining pairs are tested for co-occurrence and mutual exclusivity by Fisher’s exact test and DISCOVER. Benjamini-Hochberg *q*-values are then calculated separately within each experiment, method, and direction. Pearson correlation, which is equivalent to the phi coefficient for binary variables, is reported for each candidate pair as a descriptive effect size.

To evaluate portability, we applied the same workflow to *Pseudomonas putida* ALE mutational data from ALEdb. Only species-specific configuration values and curation tables were changed. After quality control, the *P. putida* compendium comprised 2,419 mutations affecting 402 genes across 189 endpoint samples from 46 experiments. Of these, 35 experiments contained at least 2 endpoints, and 28 contained at least 1 pair that passed the testability filter, yielding in total 460 pairs that were testable. For co-occurrence, 55 pairs from 1 experiment were testable by Fisher’s exact test, compared with 327 pairs from 24 experiments by DISCOVER. For mutual exclusivity, 20 pairs from 1 experiment were testable by Fisher’s exact test, compared with 194 pairs from 20 experiments by DISCOVER.

At a 5% false discovery rate threshold, neither method identified a significant pair. This may reflect the small size of most *P. putida* experiments: 34 of the 35 experiments contained 6 or fewer endpoints and therefore could not yield Fisher-testable pairs. Among these 34 experiments, DISCOVER yielded testable pairs in 27 experiments, but none of these pairs was significant.

## Discussion

In this study, we systematically tested gene pairs for co-occurrence and mutual exclusivity across a curated compendium of 141 *Escherichia coli* ALE experiments. To our knowledge, this is the first systematic study to formally test both patterns across such a broad collection of ALE experiments. To assess the robustness of the detected patterns to different statistical assumptions, we applied two methods, Fisher’s exact test and DISCOVER, to the same dataset. DISCOVER tested more pairs across more experiments, but only 5 pairs were significant, all for mutual exclusivity. Fisher’s exact test evaluated fewer pairs but identified 74 significant pairs, including 36 co-occurrences and 38 mutual exclusivities. Overall, co-occurrence and mutual exclusivity analysis provides a useful screening strategy for identifying candidate gene pairs for mutation-level investigation and experimental tests of epistasis.

The difference between Fisher’s exact test and DISCOVER arose primarily from the null distributions. Fisher’s exact test conditions on the observed numbers of endpoints in which each of the two genes is mutated and evaluates their overlap under a hypergeometric distribution. DISCOVER instead allows the mutation probability of each gene to vary among endpoints, accounting for heterogeneity among endpoints in the total number of mutated genes, and evaluates the overlap under a Poisson-binomial distribution [22]. Among the Fisher-significant pairs, the two methods produced similar expected overlaps, but DISCOVER’s broader null distributions assigned larger raw *p*-values to the same observed overlaps. To determine whether this difference was driven mainly by Fisher’s conditioning on fixed counts or by DISCOVER’s endpoint-specific mutation probabilities, we used a binomial model as a diagnostic comparison. The binomial model allowed the numbers of endpoints in which each gene was mutated to vary, as in DISCOVER, but retained constant mutation probabilities across endpoints. For the 74 significant pairs, its raw *p*-values were much closer to those from DISCOVER than to those from Fisher’s exact test. Together, these results support differences in fixed-margin conditioning and associated null dispersion as the main explanation for the Fisher-DISCOVER raw *p*-value differences.

Fisher’s exact test identified 74 significant pairs, while DISCOVER identified only 5. The larger set of Fisher-significant pairs, however, required careful biological interpretation. Most pairs significant only under Fisher arose in experiments with multiple strain labels or involved a gene showing single-mutation recurrence. In STE_BOP, for example, mutation patterns followed 3 endpoint groups: genes mutated in the same groups appeared to co-occur, whereas genes mutated in different groups appeared to be mutually exclusive. Similarly, single-mutation recurrence (the same distinct mutation across all affected endpoints) may reflect independent convergence [6], but it may also reflect shared ancestry or another source of dependence among endpoints. Recurrence alone does not distinguish among these explanations. Shared ancestry or other dependence among endpoints can generate significant gene-pair associations without requiring a genetic interaction between the two genes. These findings indicate that many Fisher-only results require closer examination of endpoint strain annotations, ancestral relationships, and mutation records before being prioritized for experimental validation.

The 5 pairs significant under both methods provide a useful high-priority set because their signals remained significant under both Fisher’s fixed-margin null and the greater-variance DISCOVER null. All 5 were mutually exclusive pairs. None of these 5 pairs arose from experiments with multiple strain labels, and both genes in each pair were represented by more than one distinct mutation across the affected endpoints. Significance under both methods does not establish epistasis, but it indicates that the pair is less sensitive to the choice of null model, a form of robustness that is useful for candidate prioritization, particularly when it is accompanied by a plausible biological interpretation. The 5 pairs represent potential alternative adaptive routes. In the high-temperature experiment, the mutual exclusivity pairs *ilvL*-*rho* and *mrdA*-*rho* followed the broader division between adaptation through *rho* and adaptation through RNA polymerase genes [20]. In the glycerol adaptation experiment, *crr* and *cyaA* both affect cAMP-CRP signaling and may provide alternative routes toward a similar regulatory state [26]. If the two routes provide overlapping adaptive effects, mutation of the second gene after one route has been taken may offer limited additional benefit, providing an explanation for their mutual exclusivity. The *rpoB*-*rpoC* and *rpoB*-*rpoD* pairs may also represent alternative ways to modify transcriptional regulation, since adaptive mutations in RNA polymerase components can produce broad changes in gene expression [32, 37]. These interpretations are biologically plausible, but experimental validation is still required to determine whether the corresponding mutation combinations exhibit epistasis.

The *rho-rpoB* pair illustrates directly why significance under both methods should be treated as a useful prioritization criterion rather than an absolute requirement. This pair was significant under Fisher’s exact test and fell slightly above the DISCOVER threshold, so it is not among the 5 pairs significant under both methods. Yet engineered combinations of specific recurrent *rho* and *rpoB* mutations demonstrated negative epistasis at 42.2 °C [34]. Its failure to reach the DISCOVER threshold is consistent with the broader DISCOVER null and with gene-level aggregation, which integrates the strongly exclusive recurrent *rho* mutations with other *rho* mutations that frequently occur alongside RNA polymerase mutations. This case shows that pairs significant under both methods can be prioritized as more statistically robust candidates, but biologically supported Fisher-only pairs should not be excluded without further investigation.

Most experiments in the compendium contained few endpoints, limiting their capacity for well-powered screening of co-occurrence and mutual exclusivity. In the power analysis for Fisher’s exact test, each gene was assigned a mutation probability of 0.15 per endpoint, and endpoints were assumed to be independent. A single pre-specified pair tested (without multiple-testing correction) at *p* < 0.05 required 70 endpoints to achieve 80% power for co-occurrence at OR = 10, whereas mutual exclusivity at OR = 0.1 required 215 endpoints. Only 42C_Tenaillon, which contained 111 endpoints, exceeded the 70-endpoint requirement, and no experiment reached the 215-endpoint requirement. Under a more stringent screening scenario in which many gene pairs are evaluated simultaneously, we applied fixed Bonferroni thresholds based on the numbers of Fisher-testable pairs in each direction in 42C_Tenaillon. Under this benchmark, 80% power required 231 endpoints for co-occurrence and 590 endpoints for mutual exclusivity. None of the 141 analyzed experiments reached either requirement. These values show that broad screening for gene-pair associations may require substantially more independent endpoints than are available in most current ALE experiments.

Our power analysis complements earlier efforts to guide ALE experimental design. ALEsim was used to evaluate how passage size affects adaptation and resource requirements [39], whereas our analysis addresses the number of independent endpoints needed for gene-pair association testing. Increasing the number of independently evolved lineages within an experiment may therefore be important for future ALE studies aimed at identifying convergent mutational targets and genetic interactions. Automated ALE platforms and specialized biofoundries can reduce the practical constraints of maintaining large numbers of parallel lineages and enable highly replicated evolution campaigns [40]. Greater replication can strengthen evidence for convergent mutational targets. Our power analysis further suggests that increasing the number of independent endpoints could improve the statistical power to detect co-occurrence and mutual exclusivity patterns.

The analysis pipeline was implemented as a configurable Nextflow workflow, automating gene-pair screening across experiments and enabling reproducible execution and straightforward adaptation to other species and datasets. Dataset-defining decisions, including starting-strain mutation removal, hypermutator removal, gene-name normalization, and experiment exclusion, are supplied through external tables rather than embedded in the code. The same cleaned matrices can be analyzed using Fisher’s exact test, DISCOVER, and descriptive correlation. The unit of testing can also be changed from genes to operons, pathways, or other user-defined groups. We demonstrated this portability by applying the workflow to *P. putida* from ALEdb, with changes only to species-specific configuration values and curation tables. Together, the Nextflow implementation and external configuration files support transparent and reproducible reanalysis as ALE databases expand and as alternative curation decisions or biological units are considered.

## Conclusions

This study provides a reproducible framework for screening gene-pair co-occurrence and mutual exclusivity in ALE mutation data. Comparison of Fisher’s exact test and DISCOVER assesses the sensitivity of candidate associations to null-model choice, while endpoint metadata and mutation-level evidence help interpret these associations and prioritize candidates for experimental investigation. The power analysis provides sample-size benchmarks under pre-specified scenarios and indicates that broad screening may require substantially more independent endpoints than most experiments in the compendium contain. The configurable Nextflow pipeline supports reuse across ALEdb species and user-supplied mutation datasets. This framework narrows the search space for experimental investigation of epistasis, provides benchmarks for planning future ALE experiments, and may inform combinatorial strain design.

## Methods

### Data collection

Mutation data were obtained from ALEdb (aledb.org) [17], a public database of adaptive laboratory evolution experiments and their associated mutations and experimental metadata. In addition to the public uploads, four uploads that are not public in ALEdb (STE_BOP, GLV, stALE and MS56_ALE_2022) were used with permission. When ALEdb held several uploads (duplicates) under one experiment name for a species, only one was used. The resulting data contained 286,918 mutation records from 370 experiments and 5,569 sequenced samples, across 17 organisms.

### Quality control

#### Species normalization

ALEdb contains experiments from multiple bacterial species, with species names represented by different labels. Species annotations were therefore manually normalized before species-specific filtering. The primary analysis was restricted to *Escherichia coli*, with all subspecies retained, yielding 231,441 mutation records from 269 experiments. K-12 MG1655 accounted for the largest proportion of records (109,873), with MDS42, BW25113, W3110, REL606, and other strain backgrounds also represented. The resulting compendium therefore encompassed multiple *E. coli* strain backgrounds.

*Pseudomonas putida* was selected independently using the same normalized species field to assess workflow portability, yielding 15,448 mutation records from 46 experiments.

#### Condition normalization

ALE experiments in ALEdb are conducted under diverse conditions, described with varying levels of detail and terminology across studies. Conditions were therefore curated in two stages. First, a condition was assigned to each experiment from its supplementary information, DOI and experiment name. Second, conditions with equivalent meanings but different nomenclature were normalized to a single name, so that “paraquat” and “PQ(0.8mM)” both became “Paraquat”. Experiments whose selective pressure could not be resolved from the metadata were labeled “Unclear”, and experiments serving as untreated comparison groups were labeled “Untreated”. Samples in the “Untreated” group were removed, since they are not adaptive evolution under a defined selective pressure, which excluded 1,227 mutation records from 16 experiments. All other annotated conditions were retained.

#### Mutation data cleaning

After species selection and condition annotation, a series of mutation cleaning steps was applied. The main mutation data cleaning workflow follows a previous ALE metadata analysis [6]. The number of *E. coli* mutations retained after each step is summarized in **Figure 1A**.

1. **Starting strain mutation removal**. When an experiment included a sequenced starting-strain sample, labeled A0 F0 I1 R1 in ALEdb, the mutations called in that sample were removed from every sample of the experiment, matched by reference sequence, position and sequence change.
2. **Hypermutator ALE removal**. An ALE replicate was predicted to host a hypermutator strain when both conditions were met: a coding mutation was present in a known hypermutator gene, and the replicate’s mutation count was a Tukey outlier (above Q3 + 1.5 × IQR) against the distribution of counts across all ALE replicates. The hypermutator gene list was collected from established literature. Replicates carrying 1,000 or more mutations were also removed.
3. **Mutation frequency filtering**. Only mutations reaching fixation in their sample were retained, that is, a reported presence of 1.0. This filter can be modified based on users’ needs.
4. **Endpoint definition and mutation de-duplication**. Following ALEdb, an ALE denotes one independently evolved line within an experiment, identified by its A# value. For each ALE, only the final flask was retained, preventing mutations persisting across time points from being counted repeatedly. All sequenced isolates and technical replicates from the final flask were pooled into a single endpoint, identified by experiment, ALE, and flask. A mutation was considered present if detected in any contributing sample. Of the 929 endpoints, 204 contained multiple sequenced isolates, with a maximum of 18. Within each endpoint, mutations were deduplicated by genomic position, mutation type, sequence change, mutation details, and gene annotation, leaving one entry per distinct mutation.
5. **Gene name standardization and removal of multi-gene annotations**. Gene identifiers were mapped to standard names using the protein alias table for *E. coli* K-12 MG1655 (v12.0) from the STRING database [41]. An identifier already recognized as a standard name in STRING was kept unchanged, and any other identifier was resolved by alias lookup. To prevent erroneous gene merging caused by alias collisions in STRING, primary annotations (RefSeq_gene, UniProt_GN_Name) were prioritized over synonym annotations when a name mapped to more than one entry. Annotation artifacts were normalized before lookup, stripping a trailing “CDS” and collapsing a feature annotated twice as “G|G” to “G”. Intergenic mutations were assigned to the closer flanking gene using the two distances reported in the mutation details. This resolved 3,018 intergenic entries, and the 2 whose distances could not be parsed were removed. Mutations spanning several genes, such as the amplifications and large deletions, cannot be assigned to one unit and were removed, accounting for 741 entries. A single-gene identifier that STRING could not name was retained as it is, keeping 2,173 entries that cover pseudogenes, RNA genes, and locus tags from non-MG1655 assemblies. After these steps 16,998 entries remained, across 3,733 distinct genes.
6. **Experiment exclusion and de-duplication**. Experiments were excluded if they had documented contamination or another known problem, used an engineered mutator background (Δ*mutY*, inducible *mutS*, or Δ*dnaQ*), were the wild-type controls of the Δ*mutY* series, or duplicated another deposit. In total, 25 were removed.

The resulting cleaned endpoint-level mutation data were then used to construct experiment-specific gene-by-endpoint matrices for pairwise statistical analysis.

### Pairwise association testing

#### Analysis units and matrix construction

Each experiment was analyzed separately using a binary gene-by-endpoint matrix. An entry was set to 1 if at least 1 mutation was assigned to that gene in an endpoint and 0 otherwise, so multiple mutations in the same gene and endpoint were represented as a single gene-level event. Experiments were not pooled because differences in strain background, selection pressure, and experimental duration could otherwise produce associations driven by between-experiment differences.

Within each experiment, candidate genes were required to be mutated in at least 2 endpoints, representing the minimum criterion for recurrence. Experiments with only 1 endpoint were therefore excluded because no gene could meet this criterion, leaving 141 experiments and 899 endpoints. Across experiments, 1,017 experiment-specific genes met the recurrence criterion, yielding 16,528 candidate gene pairs from 117 experiments containing at least 2 candidate genes.

#### Null models

Fisher’s exact test and DISCOVER were applied to the same experiment-specific binary matrices and candidate gene pairs. Both methods were evaluated separately for co-occurrence and mutual exclusivity using one-sided upper- and lower-tail tests, respectively. Method-specific testability and multiple-testing correction are described separately below.

Fisher’s exact test defines the null distribution conditional on the observed numbers of endpoints in which the two genes are mutated. For two genes *A* and *B*, mutated in *c*_*A*_ and *c*_*B*_ of the *N* endpoints in an experiment, let *X* denote the number of endpoints in which both genes are mutated (overlap). Under the null hypothesis of independence and conditional on these margins,

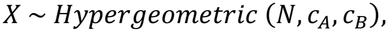

with probability mass function

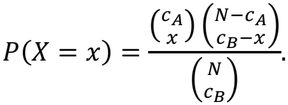

The possible overlap counts *x* range from

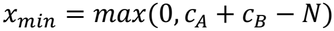

to

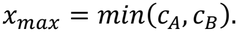

The expectation and variance are

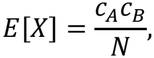

and

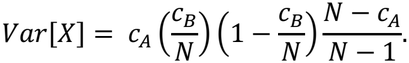

For an observed overlap *x*_*obs*_, co-occurrence was tested using the upper tail,

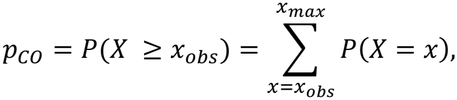

and mutual exclusivity using the lower tail,

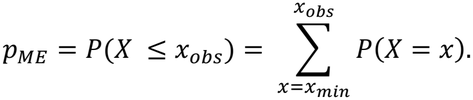

DISCOVER defines the null distribution from a fitted background probability matrix rather than by conditioning on the observed pairwise margins. For each gene in an experiment, it estimates a background probability *p*_*ij*_ that gene *i* is mutated in endpoint *j* by maximizing information entropy. The probabilities are constrained so that the expected number of endpoints in which each gene is mutated matches the observed number, and the expected number of mutated genes in each endpoint matches the observed number [22]. The background model was fitted separately for each experiment using all mutated genes, including genes that did not meet the recurrence criterion for pairwise testing. The resulting probabilities therefore account for both gene-specific mutation prevalence and differences in overall mutation burden among endpoints.

For a pair of genes *A* and *B*, let *p*_*Aj*_ and *p*_*Bj*_ denote their estimated mutation probabilities in endpoint *j*. Under the fitted independence model, their probability of overlapping in that endpoint *q*_*j*_ is:

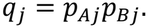

The total overlap count across *N* endpoints therefore follows a Poisson-binomial distribution,

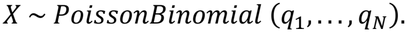

Its probability mass function is

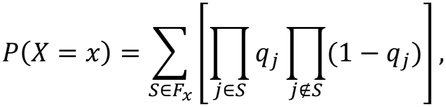

where

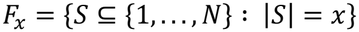

is the set of all combinations of *x* endpoints. The expectation and variance are

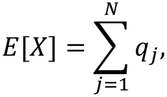

and

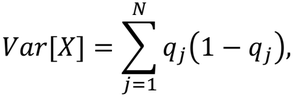

For an observed overlap *x*_*obs*_, co-occurrence was tested using the upper tail,

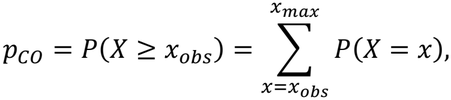

and mutual exclusivity using the lower tail,

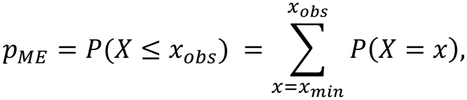

where *x*_*min*_ and *x*_*max*_ denote the minimum and maximum overlap counts allowed by the fitted Poisson-binomial null. Their determination is described in the next section.

#### Pair testability

Some candidate pairs cannot attain a raw *p*-value below 0.05 for a given direction of testing (co-occurrence or mutual exclusivity), even at the most extreme overlap allowed by the corresponding null distribution. For example, under Fisher’s exact test, if two genes are each mutated in two of four endpoints, the smallest attainable overlap is zero endpoints, representing the most extreme outcome in the mutual exclusivity direction. Even at this overlap, the one-sided *p*-value is 0.167. Thus, this pair cannot yield *p* < 0.05 for mutual exclusivity under Fisher’s exact test. An illustration of the testability filter is shown in **Figure S1**.

Testability was therefore assessed separately for each candidate pair and each direction of testing using its minimum attainable one-sided *p*-value, denoted *p*_*min*_. A pair was considered testable for a given direction when *p*_*min*_ < 0.05. Passing this filter means that a raw *p*-value below 0.05 is attainable for that pair and direction under the corresponding null distribution. It does not imply that the observed overlap yields *p* < 0.05. For a given null model, *p*_*min*_ is determined by the quantities defining that null distribution rather than by the observed overlap of the pair. For Fisher’s exact test, these are *N* and the observed numbers of endpoints in which genes *A* and *B* are mutated, *c*_*A*_ and *c*_*B*_, whereas for DISCOVER they are the fitted endpoint-specific mutation probabilities.

Pairs that could not attain this threshold were excluded from the set of tests entering the corresponding multiple-testing correction. Testability was evaluated separately for co-occurrence and mutual exclusivity because the minimum attainable *p*-value can differ between the two directions.

The attainable overlap range for a given pair depends on the null model and was determined separately for Fisher’s exact test and DISCOVER.

Under Fisher’s exact test, the observed numbers of endpoints in which the two genes are mutated, *c*_*A*_ and *c*_*B*_, are fixed. For *N* endpoints, the minimum and maximum possible overlaps are

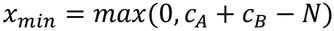

and

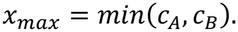

The minimum attainable *p*-value for co-occurrence is

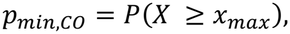

whereas that for mutual exclusivity is

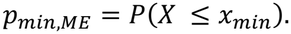

Both probabilities are evaluated under the hypergeometric null distribution.

Under DISCOVER, the overlap count follows the fitted Poisson-binomial distribution with endpoint-specific overlap probabilities

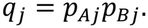

When all *q*_*j*_ lie strictly between 0 and 1, the number of overlapping endpoints can range from 0 to *N*. If *q*_*j*_ = 1 for one or more endpoints, overlap is guaranteed in those endpoints and the minimum possible overlap increases accordingly (> 0). Conversely, if *q*_*j*_ = 0 for one or more endpoints, overlap is impossible in those endpoints and the maximum possible overlap decreases (< N). The minimum and maximum attainable overlaps are therefore

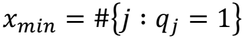

and

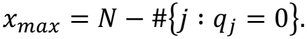

The corresponding minimum attainable *p*-value for co-occurrence is

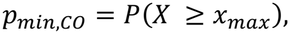

whereas that for mutual exclusivity is

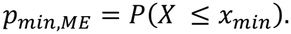

These probabilities are evaluated under the fitted Poisson-binomial null distribution.

#### Multiple-testing correction

Raw *p*-values for testable pairs were adjusted using the Benjamini-Hochberg procedure, separately within each experiment, method, and direction. Only pairs classified as testable for the corresponding method and direction were included in the adjustment. Pairs with *p*_*min*_ ≥ 0.05 were excluded from further significance testing in that direction because no overlap allowed by the corresponding null distribution could yield a raw *p*-value below 0.05 and were therefore not assigned an adjusted *p*-value. Co-occurrence and mutual exclusivity were adjusted separately because they represent distinct one-sided hypotheses and can have different sets of testable pairs. Adjustment was performed separately by experiment, consistent with the experiment-specific analysis described above, so false discovery rate control applies within each experiment, method, and direction rather than across the full compendium. Standard Benjamini-Hochberg adjustment was applied to both Fisher’s exact test and DISCOVER to ensure that differences between the two analyses reflected their null models rather than different multiple-testing procedures. Although DISCOVER provides a discrete false discovery rate procedure for discrete test statistics, this procedure was not used. Pairs with a Benjamini-Hochberg-adjusted *p*-value of *q* < 0.05 were considered significant.

### Diagnostic analysis

To assess whether the Fisher-DISCOVER difference was driven primarily by the difference in null dispersion associated with Fisher’s fixed-margin conditioning, rather than by DISCOVER’s modeling of heterogeneity among endpoints in the total number of mutated genes, a binomial model was used as a diagnostic comparison. Under the binomial model, for genes *A* and *B*, mutated in *c*_*A*_ and *c*_*B*_ of *N* endpoints, their mutation probabilities were estimated from the observed proportions of mutated endpoints,

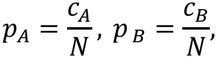

and were assumed to be constant across endpoints. Under independence, the probability *r* that both genes are mutated in an endpoint is

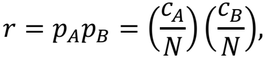

and the overlap count follows

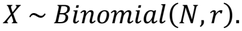

Like Fisher’s exact test, the binomial model does not introduce endpoint-specific mutation probabilities and therefore does not account for heterogeneity among endpoints in the total number of mutated genes. Its expected overlap is identical to that of Fisher’s exact test,

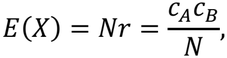

whereas its variance is

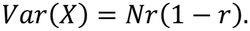

Unlike Fisher’s exact test, the binomial model does not condition on the observed numbers of endpoints in which the two genes are mutated, so these counts are allowed to vary under the null, as in DISCOVER. The binomial model therefore removes Fisher’s fixed-margin conditioning while retaining constant mutation probabilities across endpoints, providing an intermediate null for examining whether the Fisher-DISCOVER raw *p*-value differences are primarily associated with differences in conditioning or with endpoint-specific mutation probabilities.

For an observed overlap *x*_*obs*_, one-sided binomial *p*-values were calculated as

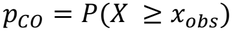

for co-occurrence and

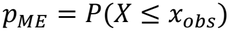

for mutual exclusivity. The binomial model was used only as a diagnostic comparison and did not define the primary set of significant gene pairs. Testability was assessed using the same *p*_*min*_ < 0.05 criterion described above. For the candidate pairs analyzed here, *r* > 0. When 0 < *r* < 1, the attainable overlap ranges from 0 to *N*, giving

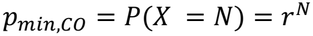

and

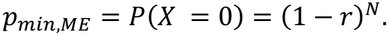

When *r* = 1, overlap is fixed at *N*, so *p*_*min*_ = 1 in both directions and the pair is not testable.

For the diagnostic significance summaries reported in the **Results**, binomial *p*-values were adjusted using the same Benjamini-Hochberg procedure as the primary analyses, separately within each experiment and direction over the pairs testable in that direction. Direct comparisons among Fisher’s exact test, the binomial model, and DISCOVER were based on raw *p*-values, so that differences in testable-set size and multiple-testing correction did not affect the comparison of their null distributions.

### Pipeline implementation

The workflow was implemented using Nextflow DSL2 and requires Nextflow version 26.04 or later. It comprises 8 processes: 6 sequential QC stages, QC report generation, and pairwise association analysis. All processes use the same Docker image built from pinned package specifications, with Conda packages identified by URL and checksum and Python packages by hash. The image contains Python 3.12, NumPy 1.26.4, pandas 2.3.3, SciPy 1.16.3, statsmodels 0.14.5, and DISCOVER 0.9.6.

QC and analysis settings are supplied through two YAML configuration files. A separate parameter file specifies the paths to these files and the input and output directories. Settings must be provided explicitly, without fallback defaults. Missing or invalid settings terminate execution with an error message identifying the affected setting. Each run writes to a separate output directory and records its parameters, container image, and Git commit identifier. Records removed at each QC stage are retained with their reasons for exclusion. With the -resume option, Nextflow can reuse cached outputs from tasks whose code, inputs, configuration, and container image remain unchanged, provided the corresponding cache records and task outputs are retained. Running the *E. coli* analysis requires Docker, more than 8 GB of memory, and approximately 1.2 GB of disk space for outputs plus 5 GB for the container image.

### Power analysis for Fisher’s exact test

Testability, as defined above, determines under a null model whether a gene pair can attain a raw *p*-value below the specified significance threshold for at least one possible overlap. Statistical power addresses a different question: the probability that Fisher’s exact test rejects the null hypothesis when an association of a specified magnitude is present. We therefore performed a power analysis for Fisher’s exact test using pre-specified effect sizes. The calculation followed the framework for Fisher’s exact test when the table margins are not fixed [42].

For two genes *A* and *B*, let *f*_*A*_ and *f*_*B*_ denote their marginal probabilities of being mutated in an endpoint. An alternative association was specified by an odds ratio *θ*. Let

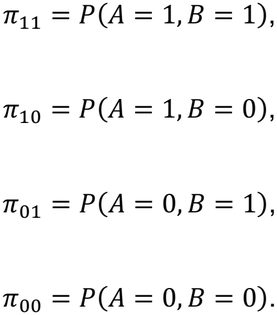

For specified values of *f*_*A*_, *f*_*B*_, and *θ*, these probabilities satisfy

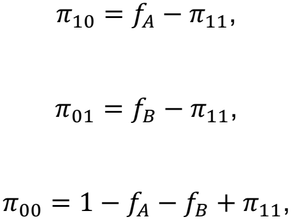

and

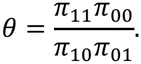

For *θ* ≠ 1, the joint probability *π*_11_ was obtained as the feasible root of

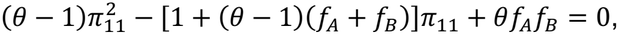

subject to

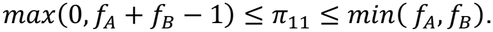

For *θ* = 1, corresponding to independence, *π*_11_ = *f*_*A*_*f*_*B*_.

The power calculation treated the *N* endpoints as independent observations generated under this specified alternative model. The numbers of endpoints in the four cells of the resulting 2 x 2 table, *n*_11_, *n*_10_, *n*_01_, *n*_00_, therefore follow a multinomial distribution,

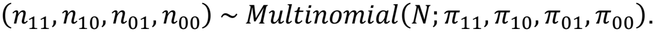

Thus, the multinomial distribution describes the possible 2 × 2 tables that could arise across repeated experiments when the marginal mutation probabilities and odds ratio are fixed at the values specified under the alternative.

For each possible 2 × 2 table with total *N*, the corresponding one-sided Fisher *p*-value was calculated using the realized margins of that table. Co-occurrence was tested using the upper hypergeometric tail and mutual exclusivity using the lower tail. A table was included in the rejection region when its one-sided Fisher *p*-value satisfied *p* < *α*. Because the margins can differ among tables generated under the alternative, the Fisher null distribution and rejection decision were evaluated separately for each possible table. Power was then calculated as the total multinomial probability of all tables in the Fisher rejection region,

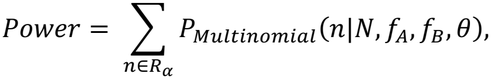

where *R* _α_ denotes the set of tables rejected by the corresponding one-sided Fisher test. The calculation was performed by exhaustive enumeration rather than Monte-Carlo simulation. For a given *N*, significance threshold, and testing direction, the Fisher rejection decision for each possible table does not depend on the alternative odds ratio, allowing the rejection region to be reused when evaluating different effect sizes.

Power was evaluated for Fisher’s exact test only. Fisher’s test provides a finite rejection region over the possible 2 × 2 tables for a given *N*, allowing power to be calculated directly under the specified multinomial alternative. An analogous calculation was not performed for DISCOVER because its null distribution is determined by an experiment-specific fitted background probability matrix. Defining an alternative for a single gene pair would therefore additionally require specifying how the full background matrix and its fitted endpoint-specific probabilities would change under that alternative.

For the reference power curves in **Figure S2**, both genes were assumed to be mutated in 15% of endpoints,

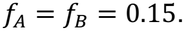

This value represents the empirical center of the marginal mutation frequencies among pairs testable by Fisher’s exact test in at least one direction, for which the pair-weighted mean was 0.149. Co-occurrence was evaluated at *θ* = 10, and mutual exclusivity at the reciprocal effect size, *θ* = 0.1. These odds ratios represent pre-specified effect sizes and were not estimated from the observed gene pairs.

Two significance thresholds were considered. For a single pair specified in advance, *α* = 0.05 was used. A second analysis provided a conservative multiple-testing benchmark based on 42C_Tenaillon, the experiment with the largest number of endpoints. Because co-occurrence and mutual exclusivity were tested and corrected separately in the primary analysis, direction-specific numbers of Fisher-testable pairs were used. These were 5,482 pairs for co-occurrence and 236 pairs for mutual exclusivity, giving fixed Bonferroni thresholds of

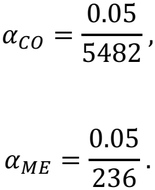

These thresholds were used as conservative fixed-multiplicity benchmarks and were not intended to reproduce the Benjamini-Hochberg procedure used for significance testing in the primary analysis.

For each reference scenario, power was calculated over increasing values of *N*. Because Fisher’s exact test is discrete, power can change non-smoothly as *N* increases. The reported sample-size requirement for 80% power was therefore defined as the smallest *N* from which power remained at or above 80% for all larger evaluated values of *N*. **Figure S2C** additionally evaluates the single-pair scenario at *α* = 0.05 across a range of pre-specified odds ratios while retaining *f*_*A*_ = *f*_*B*_ = 0.15, illustrating how power depends jointly on effect size and the number of endpoints.

For the Bonferroni benchmark, the numbers of tests were held fixed at the values observed in 42C_Tenaillon while *N* was varied. These curves therefore represent power under a fixed multiple-testing requirement. They do not model changes in the number of candidate or testable pairs that could occur as additional endpoints are added. Consequently, the resulting values should be interpreted as reference design benchmarks rather than universal sample-size requirements.

### Use of artificial intelligence tools

Claude Code (Anthropic, Opus 5.5) was used to assist with code development, and ChatGPT (OpenAI, GPT-6.1) was used to improve the language and readability of the manuscript. The authors reviewed and revised the AI-assisted code and text and took full responsibility for the accuracy and integrity of the computational analyses, interpretation of results, and final manuscript.

## Abbreviations

ALE: Adaptive Laboratory Evolution
OR: Odds Ratio
*E. coli*: *Escherichia coli*
*P. putida*: *Pseudomonas putida*

## Declarations

### Ethics approval and consent to participate

Not applicable

### Consent for publication

Not applicable

### Availability of data and materials

The analysis pipeline with its input data (https://doi.org/10.5281/zenodo.24175868) and the figure code with the processed data (https://doi.org/10.5281/zenodo.24175860) are archived on Zenodo. Both are also available on GitHub at https://github.com/Multiomics-Analytics-Group/ale-cooccurrence-mutual-exclusivity-nf and https://github.com/Multiomics-Analytics-Group/ale-cooccurrence-figures. Mutation records and experimental metadata were obtained from ALEdb (https://aledb.org), and gene aliases from STRING v12.0 (https://string-db.org). The mutation records of 4 experiments (STE_BOP, GLV, stALE and MS56_ALE_2022) are non-public datasets in ALEdb. The Zenodo archives contain only derived tables from these experiments. Detailed mutation records of these 4 experiments are available from the corresponding authors on reasonable request and with the permission of their owners.

- Project name: ale-cooccurrence-mutual-exclusivity-nf
- Project home page: https://github.com/Multiomics-Analytics-Group/ale-cooccurrence-mutual-exclusivity-nf
- Archived version: v1.0.0; https://doi.org/10.5281/zenodo.24175868
- Operating systems: Linux and macOS
- Programming languages: Nextflow DSL2, Python, and Bash
- Other requirements: Nextflow 26.04 or later and Docker. The figure code uses the conda environments in its repository.
- License: MIT. Included third-party code and data keep their own licenses.
- Restrictions on use by non-academics: None for the code. Building the Docker image downloads packages from Anaconda’s default channel, whose terms require a paid plan for for-profit organizations with more than 200 employees.

## Authors’ contributions

Conceptualization: PVP., AS., AF., BS.; Implementation: BS., AS.; Data processing and generation: BS.; Writing: BS., PVP., AS., AF.; Supervision: AS., PVP., AF. All authors read and approved the final manuscript.

## Acknowledgements

The authors gratefully acknowledge Bernhard Palsson, Lei Yang, Emre Ozdemir, Xuwen (Amy) Lou, and Gaoyuan Li for their valuable discussions and Josefin Johnsen for her laboratory work on the MS56_ALE_2022 experiment.

## Author Information

## Corresponding Author

**Alberto Santos**: The Novo Nordisk Foundation Biotechnology Research Institute for the Green Transition, Technical University of Denmark, 2800 Kongens Lyngby, Denmark; Department of Health Technology, Technical University of Denmark, Kongens Lyngby, Denmark

**Patrick Victor Phaneuf**: The Novo Nordisk Foundation Biotechnology Research Institute for the Green Transition, Technical University of Denmark, 2800 Kongens Lyngby, Denmark

## Supplements

**Figure S1.**
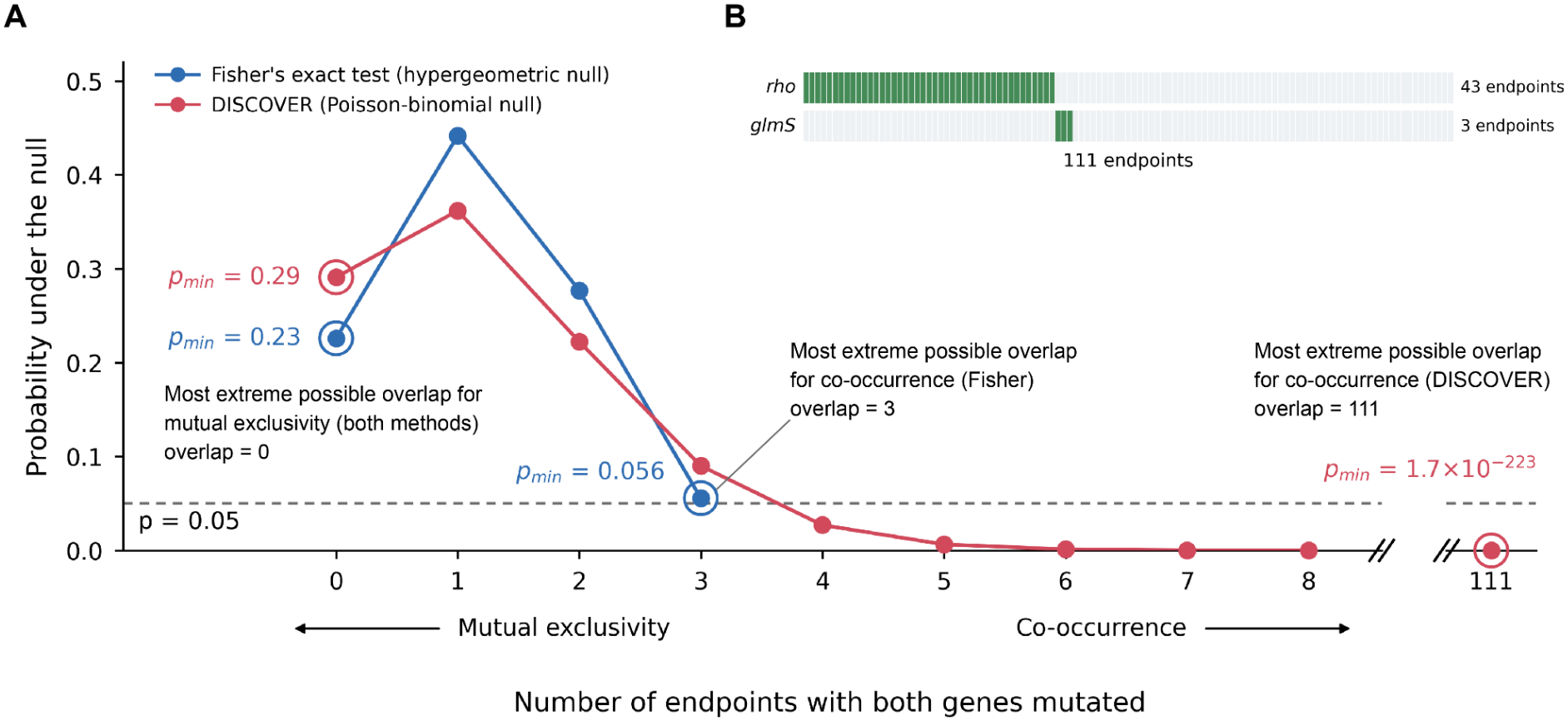
Illustration of gene-pair testability under Fisher’s exact test and DISCOVER. **(A)** Null distributions of overlap (the number of endpoints in which both genes are mutated) for *glmS* and *rho* in the 42C_Tenaillon experiment under Fisher’s exact test (blue, hypergeometric null) and DISCOVER (red, Poisson-binomial null). Circled points mark the minimum and maximum attainable overlaps under each null model. The probability at each circled point equals the minimum attainable one-sided *p*-value, *p*_*min*_, for the corresponding method and direction. A pair passes the testability filter when *p*_*min*_ < 0.05 (dashed line). For mutual exclusivity, the minimum overlap is 0 under both methods, giving *p*_*min*_ = 0.23 (Fisher) and 0.29 (DISCOVER). The pair is therefore untestable for mutual exclusivity despite zero observed overlap. For co-occurrence, Fisher’s exact test fixes the observed numbers of endpoints in which each gene is mutated, limiting the maximum overlap to 3 and giving *p*_*min*_ = 0.056. Under the fitted DISCOVER null, these numbers are not fixed, allowing a maximum overlap of 111 and giving *p*_*min*_ = 1.7 x 10^-223^. The pair is therefore testable only for co-occurrence under DISCOVER. The x-axis is broken between 8 and 111. **(B)** Observed mutation patterns across 111 endpoints. *rho* is mutated in 43 endpoints and *glmS* in 3, with no overlap. Green indicates mutation presence and light gray absence.

**Figure S2.**
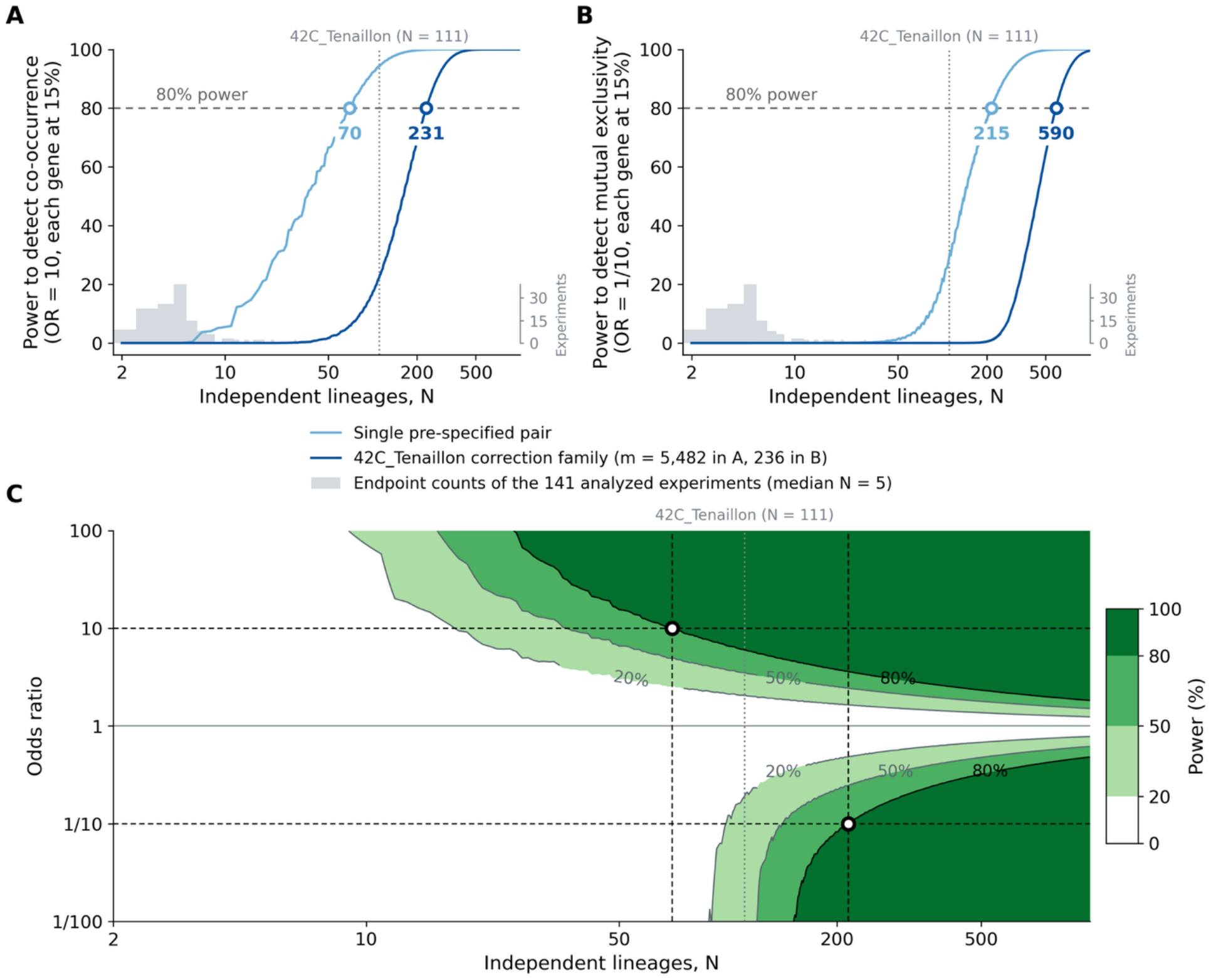
Power to detect gene-pair associations depends on lineage number, multiple-testing burden, and association direction. Power of one-sided Fisher’s exact test was evaluated for pre-specified effect sizes, assuming independent lineages and a marginal mutation probability of 0.15 for each gene. This probability matches the empirical center of marginal mutation frequencies among pairs testable by Fisher’s exact test in at least one direction (mean, 0.149). (**A**) Power to detect co-occurrence at OR = 10 as the number of independent lineages (*N*) increases. (**B**) Power to detect mutual exclusivity at OR = 0.1. Light-blue curves show testing of a single pre-specified pair (no FDR correction) at *p* < 0.05; dark-blue curves use fixed Bonferroni thresholds of *p* < 0.05/*m*, with *m* = 5,482 for co-occurrence and 236 for mutual exclusivity, based on Fisher-testable pair counts in 42C_Tenaillon. Open circles and adjacent labels mark the smallest *N* from which power remains at or above 80% across larger evaluated sample sizes: 70 and 215 lineages for single-pair testing, and 231 and 590 under the Bonferroni benchmarks, respectively. Gray bars show the distribution of endpoint counts across the 141 analyzed experiments (median, 5; maximum, 111). (**C**) Single-pair power at *p* < 0.05 across lineage number and pre-specified odds ratio. Odds ratios above and below 1 are evaluated using the upper and lower tails of Fisher’s exact test, respectively. Shading and contours indicate 20%, 50%, and 80% power. The horizontal dashed lines mark OR = 10 and OR = 0.1. Black vertical dashed lines and open circles mark *N* = 70 and *N* = 215. Gray dotted vertical lines mark *N* = 111 in all panels.

